# Slowing down: A macroevolutionary approach to the hypometabolic strategies of amphibians

**DOI:** 10.1101/2025.07.19.665691

**Authors:** Leticia M. Ochoa-Ochoa, Juan D. Vásquez-Restrepo, Mylena Masache, Rebecca D Tarvin

## Abstract

The ability to survive harsh environmental conditions has probably been a key factor in the evolutionary success of organisms that cannot migrate long distances, such as amphibians. We expect that having a hypometabolic strategy (HS) —aestivation or hibernation— to deal with severe climates, would be a plesiomorphic trait. We 1) inferred the ancestral state of a HS, using two phylogenies for amphibians, 2) tested if species with a HS have larger distributional ranges, and 3) explored how a HS may affect amphibian assemblage resilience using multiple models of climate change. Ancestral state reconstruction for the most recent common ancestor (MRCA) of Class Amphibia showed ∼50% probability of a HS. The probability was higher for the MRCA of each Order (>70%), suggesting a widespread HS in the ancestors of modern amphibians. Phylogenetic regressions showed no relation between the probability of having a HS and the distribution range size. Climate analyses predict that tropical zones will have the greatest change in climate, involving novel harsh seasonality. Since tropical amphibian assemblages have the lowest proportion of species with HS, they may be more vulnerable to climate change. It is probable that HS have been key for the evolutionary success in amphibians, and they will likely impact their future survival in the face of climate change. Despite the potential importance of the HS for amphibians, information was available for a diverse but only a small subset of species; we urge researchers to report data on aestivation or hibernation in amphibians to facilitate future studies.

## Introduction

Most animals seek shelter when environmental conditions turn harsh. For instance, amphibians use shelters to hide during their inactive hours on a daily basis (Wells, 2007) in order to hide from predators and conserve water. Amphibians are required to keep their skin moist because it is crucial for breathing, and, therefore, their evaporative water loss is normally high (Lillywhite, 1975). Consequently, to survive long periods (weeks, months, or years) during adverse environmental conditions, amphibians resort to a type of dormancy or latency in a state of reduced or depressed metabolism (hereafter, hypometabolism) when they hide underground or underwater. The lengths of these hypometabolic periods vary considerably across species and populations because it seems to be driven by the local environmental conditions (Figure 1).

**Figure 1.**
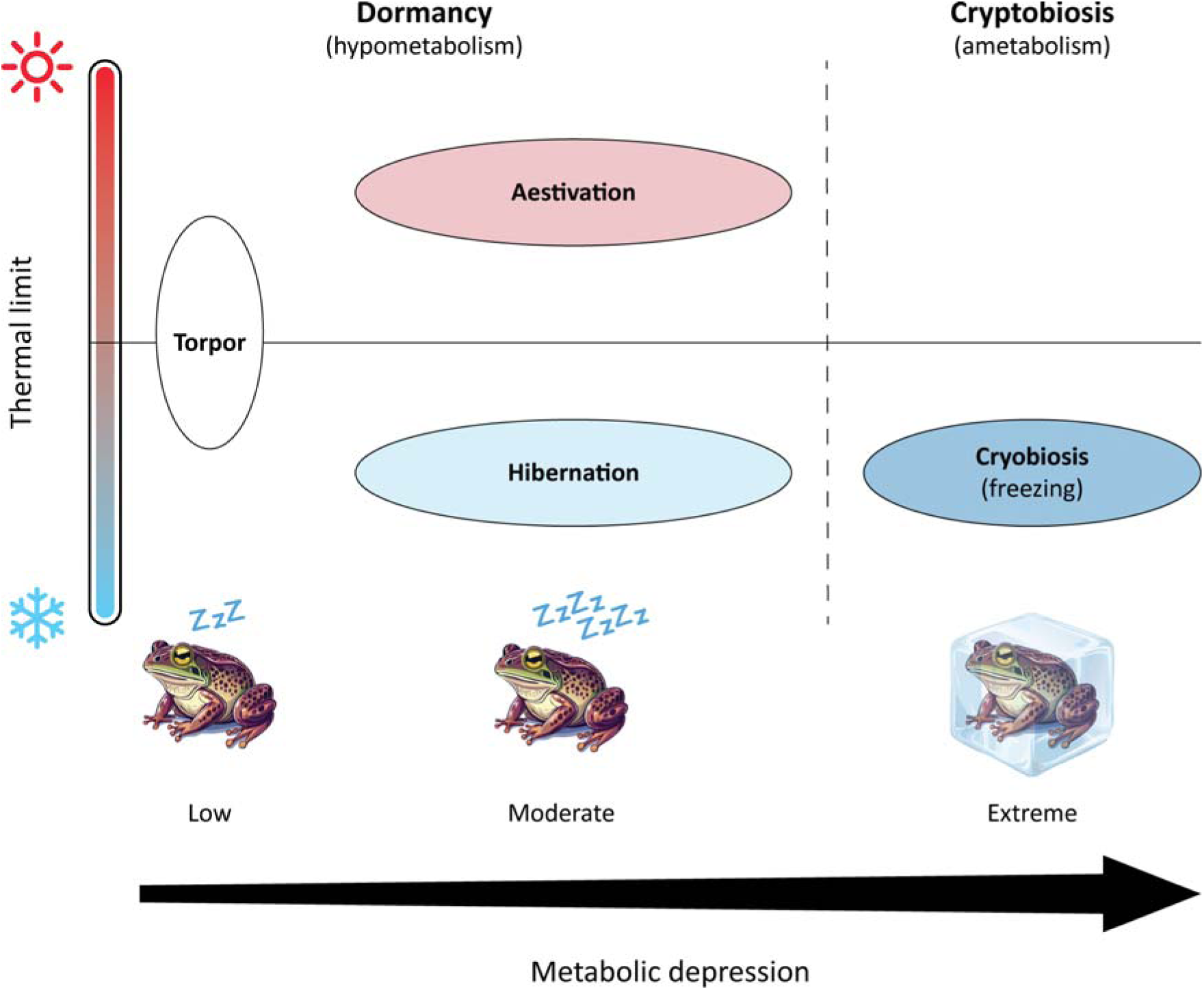
Representation of the different levels of metabolic depressions currently reported in amphibians. The ovals represent the range in which each state would be denominated as such. For example, there would be a wide range of aestivation and hibernation levels. Illustrations were generated using the Adobe Illustrator generative AI.

Here we will use the most common terminology for the different states of dormancy: we will refer to summer dormancy as aestivation and winter dormancy as hibernation (see Box 1). We also make the assumption that both strategies imply a hypometabolic state. Thus, we will refer to them collectively as hypometabolic strategies (HS). Here we will not include cryptobiosis (from the Greek *crypto* and *bios* meaning ‘hidden life’), an extreme form of hypometabolic state, where metabolism is reduced to an undetectable but non-zero level (e.g., cryobiosis, when organisms freeze in order to survive the winter), although it might be considered an extreme case of hypometabolism (Figure 1). Nonetheless, whether in the hottest and driest season of the year (i.e., aestivation) or in the coldest (i.e., hibernation), the end result is the same, some amphibians “slow their lives” (namely, lower their metabolism) in order to get through the unfavorable conditions that the environment imposes. In some cases, this pattern is so extreme that amphibians are active only a few weeks each year, like some desert amphibians (McClanahan, 1972; Cowan & Storey, 1999; Withers & Cooper, 2010). For example, the maximum aestivation period recorded in amphibians is five years in the Australian striped burrowing frog *Cyclorana platycephala* (Van Beurden, 1980). It is unknown if these extremely long periods of hypometabolism can be repeated several times throughout the animal’s life or if there are limits. In some cases, aestivating amphibians produce a cocoon from several layers of shed skin (Withers, 1995); the cocoon provides extra insulation to prevent water loss and might increase the duration of aestivation periods in these species (Withers, 1998). For example, estimations show that it would take more than eight years for a *Cyclorana* frog with a cocoon and an empty bladder to lose 30% of its body mass by evaporation (Sadowski-Fugitt *et al*., 2012).

### Box 1. The use of different terms of dormancy.

Hypometabolism is a general phenomenon that is not unique to amphibians and includes a wide range of metabolic depressions such as hibernation, aestivation, dormancy, brumation and torpor. In general, dormancy is considered to be a term that encompasses all hypometabolic states, and the key aspect is that dormancy is preventive and anticipates stressful physiological conditions to avoid harmful changes in physiology (Navas & Carvalho, 2010; Storey & Storey, 2012; Reynolds, 2019). Torpor, from the Latin *torp*ō*r* (Borror, 1960), means to be numb, inactive, or dull, and is referred to as a daily rest; it was commonly used as synonym of hibernation (i.e., winter torpor; Pinder *et al*., 1992; Boutilier *et al*., 1997; Withers & Cooper, 2010; Wilkinson *et al*., 2017), yet it should not be considered a sustained hypometabolic state such as aestivation or hibernation (Castanho & De Luca, 2001; Navas & Carvalho, 2010 but see Geiser, 2020, 2021). Aestivation or estivation, both derived from the Latin *aest*ā*s* that mean “pertaining to summer” (Borror, 1960), is defined as a state of reduced metabolism occurring in hot and dry conditions. Nevertheless, Pinder et al. (1992) consider that aestivation occurs in both hot and cold conditions and that its main driver is the lack of water. For example, toads of the genus *Scaphiopus* aestivate during winter (Seymour, 1973). Aestivation can occur when the amphibian burrows itself into the soil or seeks shelter in crevices or below logs and rocks, but it can also occur out in the open either on land or on plants, although it is much less common (Navas & Carvalho, 2010). Finally, brumation and hibernation both come from Latin words meaning “related to winter” (Borror, 1960), the former from *bruma* and the latter from *h*ī*bern*. The use of the terms brumation and hibernation may be sometimes ambiguous, as they refer to the lethargic state that some animals assume during cold conditions. However, brumation was proposed to refer to obligate hibernators that cannot control their body temperature during winter dormancy rather than those that can (Mayhew, 1965).

According to Storey & Storey (2010), three requirements must be satisfied in order for a state to be considered hypometabolic: 1) an overall strong suppression of metabolic rate (at least a 70–80% reduction); 2) differential control over the rates of various metabolic processes so that energy use is reprioritized to favor core vital cell functions; and 3) implementation of actions that protect cells and preserve viability over long periods of time. There are several mechanisms that are activated during metabolic rate depression, and it is proposed that these mechanisms, their interactions, and their regulatory signals, form a common molecular basis for all states of metabolic depression including aestivation, hibernation, and some types of anoxybiosis (metabolic depression due to a lack of oxygen; Storey & Storey, 2004). Although the relationship between environmental condition and hypometabolic state is well documented (see below), we still lack assessment of this idea in amphibians across a large taxonomic scale.

The ability to tolerate unfavorable or extremely unfavorable environmental conditions (such as very high or very low temperatures or droughts) has probably been a key factor in the evolutionary success of organisms that cannot escape such harsh conditions. As a group, amphibians have been on Earth for more than 300 Myr and there have been many periods of unfavorable conditions during that long time (Vitt & Caldwell, 2014; Jetz & Pyron, 2018; Judd *et al*., 2024), including several mass extinctions. The ability to sustain a hypometabolic state might have also been an important factor in the colonization of land, given that lungfish also aestivate and produce cocoons (Jiang *et al*., 2023). In this paper we first aim to infer the ancestral state and evolutionary trajectory of hypometabolic strategies in all living amphibians. Our approach is based on the observation that many groups of frogs with aestivation evolved in arid or semiarid environments, such as Ceratophryidae (Faivovich *et al*., 2014). All ceratophryids are capable of prolonged aestivation and cocoon formation, even the ones that no longer live in seasonal environments (Zimicz *et al*., 2024). Paleozoic amphibians appear to have burrowed in response to seasonal droughts (Hembree *et al*., 2004), and likely other climate crises. For example, radiation of fossorial amphibians and increase in abundance and complexity of burrows coincide with the Cisuralian aridification and end-Permian extinction events (Marchetti *et al*., 2024). The ability of early Tertiary pelobatids to avoid drought by estivating in burrows is thought to be a preadaptation to survive the global cooling of the middle Eocene (Henrici & Haynes, 2006). Therefore, we would expect that hypometabolic strategies evolved or were retained in places with extreme environmental conditions, specifically with aestivation occurring in semiarid conditions and hibernation in cold conditions (Hembree *et al*., 2004; Henrici & Haynes, 2006; Faivovich *et al*., 2014; Marchetti *et al*., 2024; Zimicz *et al*., 2024). As part of this aim, we then inferred the ancestral state and evolutionary trajectory of amphibian HS; given the role of amphibians colonizing land and inhabiting harsh or seasonal environments, we expected to find that the capacity for hypometabolism is a plesiomorphic trait—an ancestral trait inherited from a common ancestor and shared by all members of a clade that in some lineages may be lost subsequently— in amphibians.

As a second aim, we explore the association between the presence of hypometabolic strategies and distributional ranges. We suggest that if a species has the possibility to avoid stressful environments, it should be able to invade novel areas without negative fitness consequences. Moreover, having a hypometabolic strategy may allow a species to survive over a broad range of environmental conditions. Therefore, we expect that species with a hypometabolic strategy will have larger distribution ranges.

Finally, the presence or absence of a hypometabolic strategy has possible implications for how amphibians might respond to future changes in climate, and thus may be a concern for conservation biologists. Indeed, climate change will impact amphibian distributions and ranges (Ochoa-Ochoa *et al*., 2012; Mokhatla *et al*., 2015; Velasco *et al*., 2021), will vary in speed across regions (Ochoa-Ochoa & Velasco, 2024; García-Rodríguez et al., 2025), and produce climatic debt—a disequilibrium between species communities’ climatic tolerances and the magnitude of climatic change (He *et al*., 2023; Smith *et al*., 2025). As a third aim, we explore the vulnerability of amphibian assemblages due to climate change in relation to the presence of hypometabolic strategies. Species that can enter dormancy or a sustained hypometabolic state may be able to deal better with environmental changes such as desertification, severe temperature/precipitation changes or extreme cold conditions. Therefore, we propose that climatic resilience—the ability to cope with climatic stressors—of amphibian assemblages increases with increasing proportions of species with hypometabolism (Figure 2). Based on this hypothesis we make predictions regarding amphibian assemblage vulnerability across the globe. Amphibian hypometabolic strategies remain an understudied phenotype and here we take a macroevolutionary perspective to improve our current state of knowledge.

**Figure 2.**
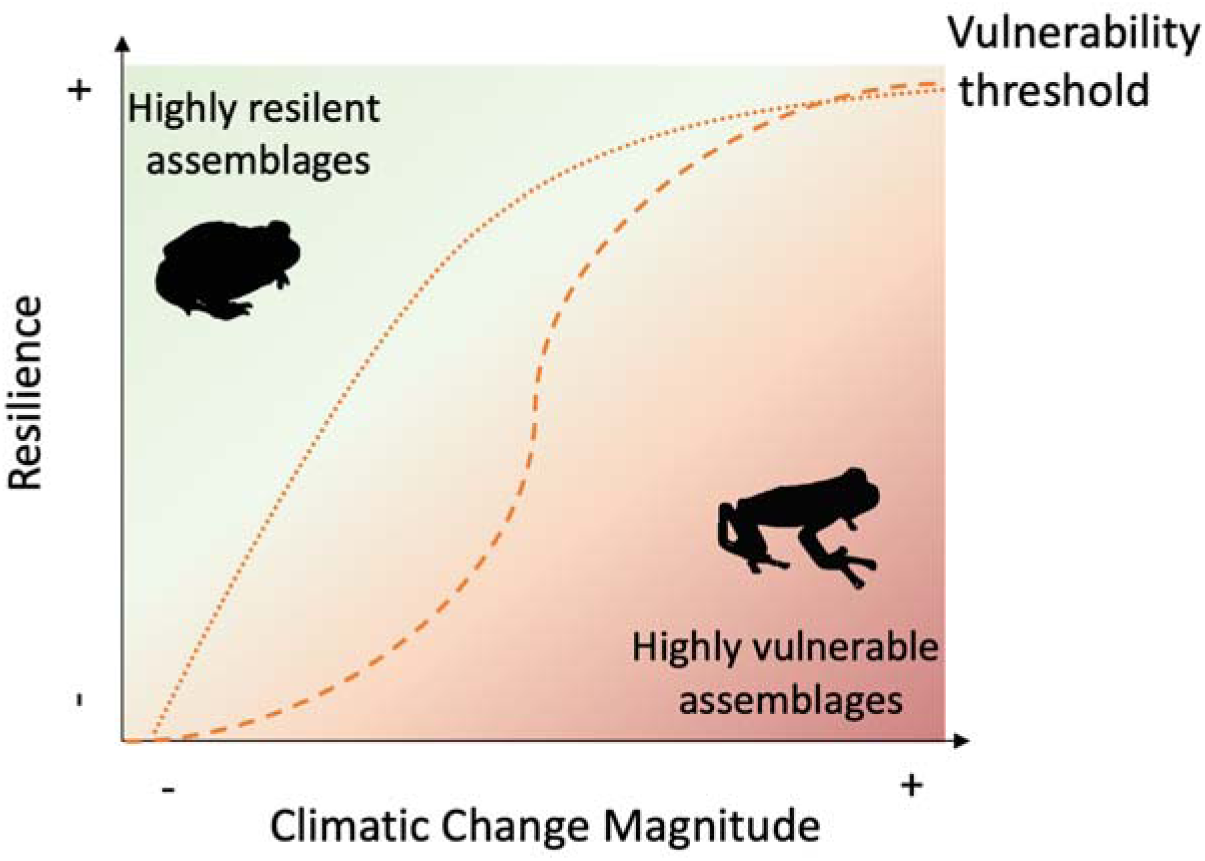
Resilience of amphibian assemblages to cope with climate change. Highly resilient assemblages are expected to have a high proportion of species with presence of hypometabolism. In contrast, highly vulnerable assemblages would be where very few species have hypometabolism. Hypothetical lines are drawn to represent two types of vulnerability transitions: abrupt threshold (dashed) and elastic threshold (dotted).

## Methods

### Hypometabolism data collection

We compiled information about hypometabolic strategies, particularly aestivation and hibernation, through a systematic literature review (see *Systematic literature review*). A comparative dataset for hypometabolic states was assembled including aestivation/hibernation status (present/possible/absent), and range size in km^2^ (Table S1 in the Supplementary Information). For scoring aestivation/hibernation, we considered them as present in a species if it was reported in at least one population or subspecies in the literature. We recorded absences (tentatively) when published studies or researchers noted its absence from personal experiences in fieldwork. However, we note that these cases will likely require further validation, and therefore, we did not include them as part of the training dataset (see *HS probability*) for subsequent models. Rather, an estimated probabilistic value based on a model was used instead.

### Systematic literature review

In order to compile the dataset of amphibians that present a hypometabolic state, an exhaustive bibliographic search was carried out. The systematic search was performed using the advanced search tool of each of the following search engines: Scopus, PubMed, and Web of Science, until December 2024. We used the words (aestivat* AND amphibian*) and (estivat* AND amphibian*) for aestivation, and (hibernat* AND amphibian*) for hibernation. Studies were selected according to the following criteria: 1) original and review manuscripts including not just peer-reviewed literature, but also theses; 2) original articles in which some biological, ecological, physiological, or genetic aspect of the aestivation process were mentioned; and 3) manuscripts where the focus was not related to dormancy processes, but an aestivating or hibernating species was mentioned. We also performed a search on AmphibiaWeb to review and complete our resulting list of species that aestivate, hibernate, or have a cryptobiotic state. Scientific names follow AmphibiaWeb (2024) nomenclature.

### Phylogenetic and geographic data

Because our subsequent analyses require a phylogenetic framework, we used the amphibian supertrees featured in Jetz and Pyron (2018) and in Portik *et al*. (2023), although these last ones are just for the Anura clade. Given that the first set of trees were generated with a mixed methodology incorporating a backbone for the species with available genetic information and the imputation of the species without data, the final consensus is highly polytomic. Therefore, in order to account for the inherent phylogenetic uncertainty (Rangel *et al*., 2015), we subsampled 100 randomly resolved trees from each dataset. In addition, we downloaded the available distribution polygons for amphibians from the International Union for Conservation of Nature Red List (IUCN, 2024; https://www.iucnredlist.org) in its 2023-1 update. Because the trees and polygons may differ in the taxonomic treatment of some species, we harmonized the names between data sources to maximize the number of species in the final dataset. To do so, we manually validated the non-matching species against their synonym list available in the Amphibian Species of the World (https://amphibiansoftheworld.amnh.org) looking for alternative names. From the 857 species with hypometabolism data, 774 from Jetz and Pyron’s dataset (approx. 90%) and 477 from Portik *et al*.’s dataset (83.52%) were shared between the phylogeny and geographic data. Although the non-matching species were not included in the phylogenetic comparative analyses, they were included to generate the hypometabolic probabilities (see *HS probability*) and to illustrate the summary information on the global status of hypometabolism in amphibians.

### HS probability

Since we have literature information for only 857 species out of the 8,823 amphibians currently described, fewer than 10% of the total number of species (AmphibiaWeb, 2024), an important issue in this study is the large amount of missing data. Thus, we calculated the probabilities of the presence of a hypometabolic strategy in the tips with unknown states by assuming that hypometabolic states are expressed in response to environmental conditions and that species are adapted to the environments they inhabit (Hembree *et al*., 2004; Henrici & Haynes, 2006; Faivovich *et al*., 2014; Zimicz *et al*., 2024). Importantly, we highlight that this approach makes the assumption that species have tracked their niches over evolutionary time. To conduct this analysis, we first extracted for each species the maximum values for the 19 WorldClim bioclimatic variables (https://www.worldclim.org) at 10 arcmin resolution (approx. 18.5 km at the equator) based on 7,949 distribution polygons. To address multicollinearity, we thinned our dataset by excluding highly correlated variables using the Variance Inflation Factor (VIF) with the *vifstep* function as implemented in ‘usdm’ v.2.1 (Naimi *et al*., 2014). We then fit a One-Class Support Vector Machine (OC-SVM) model using a subsampled training dataset that included only species with confirmed presence separately for aestivation and hibernation.

OC-SVM is suitable for presence-only data, allowing us to avoid model biasing given the uncertainty in the absence data (false negatives). For this, we employed the *svm* function in the ‘e1071’ v.1.7-16 package (Meyer *et al*., 2024), using a radial kernel with a custom gamma (shape of the kernel) for each trait (aestivation = 0.25, hibernation = 0.15), nu = 0.1 (sensitivity to outliers), and a 10 k-fold cross-validation strategy. We manually tuned the shape of the kernel to optimize the model’s performance by minimizing the number of false negatives (true presences with low estimated probability) while increasing the model’s accuracy (Figures S1–S2). Posteriorly, decision values were scaled between 0 and 1 and transformed into probabilities using a sigmoid function. Subsequently, for species with confirmed presence we kept the probability as 1, but replaced the other ones in our literature review (i.e., possible state of aestivation or hibernation) with their inferred probability in order to account for the uncertainty in defining true absences. We repeated this estimation twice, once for aestivation and once for hibernation, for each phylogeny since the number of species in the model change, thus will the probabilities. For the species assigned by the literature review as “possible”, two outcomes were possible: 1) they were assigned a low probability, equivalent to an absence; 2) they were assigned a high probability (> 80%), equivalent to a presence under our assumption that climate is triggering such hypometabolic strategies (and our initial coding from the literature is just a reflection of the absence of data).

Afterwards, in order to use these probabilities in the combined hypometabolism analysis, we assigned the maximum probability of either aestivation or hibernation to each species (i.e., if the species is predicted to estivate but not hibernate it would still be counted as a “presence” for having a hypometabolic state). Although we calculated the missing data for all the amphibians in the trees using probabilities, we performed subsequent ancestral state reconstructions (see *Reconstruction of ancestral states*) in trees with twice the number of species with hypometabolic data, by using the 774 species from the literature (fixed species) and randomly subsampling another 774 species for each tree with Jetz and Pyron (2018) phylogeny and 477 species for Portik *et al*. (2023). Thus including 50% of species with literature hypometabolic state and 50% of species with inferred hypometabolic state. The aim of this last is: 1) to increase the taxonomic representativeness of the different amphibian taxa; 2) to reduce the bias in the results by accounting only for species with presence or potential presence data; and, finally, 3) to balance the effect of using the entire estimated dataset.

### Reconstruction of ancestral states

Here, we modeled the evolutionary transitions between the presence and inferred absence of aestivation, hibernation, and hypometabolism (either aestivation or hibernation), the last following the assumption that all states of metabolic depression have a common molecular basis (Storey & Storey, 2004) but we stress that the environmental cues to enter aestivation or hibernation are very likely distinct. Subsequent analyses were performed using ‘phytools’ v.2.3 (Revell, 2024), and a script can be found as part of the Supplementary Files. First, we employed the Markov k model (hereafter Mk model) for discrete character evolution of characters with k unordered states (Lewis, 2001) to find the best state transition model, as implemented in the function *fitMk*. As a Markovian model, it assumes that transitions between states depend only on the current state and not on previous ones. We tested two different models of transition rates: equal (ER) and all different (ARD), allowing the π parameter to estimate the prior distribution of the root state jointly with the other model parameters rather than defining it *a priori* (Fitzjohn *et al*., 2009). Since we are using a collection of trees rather than a single one, we ran the analyses 100 times and chose the best model based on its likelihood and Akaike information criterion (AIC) values. For both phylogenies the best transition model for aestivation and hypometabolism was inferred to be the ARD, while for hibernation, there were no differences between the ARD and the ER (Figures S3–S4). Thus, the reconstructions were estimated using the ARD models.

Given that the presence of a hypometabolic state for many species may be ambiguous given the paucity of data, we accounted for uncertainty in trait states as probabilities. Then, we mapped possible scenarios of trait evolution using stochastic character mapping over the 100 trees for aestivation, hibernation, and hypometabolism separately. We employed the *make.simmap* function with 100 simulations per tree using the ARD transition model, π as in the fitMk, a non-fixed transition matrix (Q) estimated with a Markov Chain Monte Carlo (MCMC) approach, empirical priors from observed data, a variance of 0.1 (vQ), and a sampling frequency of 10. Thus, we ran a total 1000 MCMC iterations, sampling every 10 for a total of 100 independent histories per tree. Subsequently, we calculated the average number of transitions across 100 simulations for each of the 100 trees using the countSimmap function. Since the Q matrices were estimated independently for each simulation and tree, the MCMC algorithm may get stuck in suboptimal peaks, yielding extremely low or high numbers of transitions in some cases and increasing the uncertainty in the transition counts. Therefore, the average number of gains and losses was calculated excluding outlier values (those above or below 1.5 times the interquartile range).

Because each tree differs in topology, it is not possible to summarize the nodes across the entire set of reconstructions (10,000 different sampled histories). Thus, for the purpose of visualizing the results, we selected one of the 100 trees. Instead of choosing a random tree, we selected the one whose likelihood in the evaluation of the evolutionary models was closest to the median of the distribution frequency of all trees (Figures S3–S4). These particular trees were only used for graphical purposes, the results are based on 100 trees from each phylogeny. It is important to highlight that for the Jetz and Pyron phylogeny, given the method used for inferring the base topology, such differences (i.e., polytomies) are expected to be more pervasive at lower taxonomic levels (e.g., species or genus). Thus, to overcome topological differences while accounting for uncertainty, we identified the MRCA for superfamilies and orders, for both set of analyses, using the fixed species on each tree (i.e., those non-randomly selected), and averaged the posterior probabilities of those nodes. It is important to note that, those MRCA are not necessarily the same in all the trees but the most inclusive monophyletic groups encompassing our fixed species. Because this kind of reconstruction requires discrete traits at tips, we considered values equal or greater than 80% as strong evidence for presence, given that the probability for all true presences from the literature was estimated with the same threshold (see Figures S1 and S2). Moreover, in order to explore the transition of the states in the nodes along time, we also performed a joint ancestral state reconstruction over the 100 random trees from each phylogeny using the function *ancr*, to quantify the proportion of nodes with aestivation or hibernation on 5-Myr bins and its possible association with the mean Earth surface temperature (Judd *et al*., 2024).

### Geographic distribution of species with hypometabolism

To explore geographic patterns in our data, we generated maps of species richness with aestivation, hibernation, and hypometabolism by overlapping the IUCN species polygons (IUCN, 2024) with a grid of 1° × 1° (ca. 111 km at the equator); we also generated maps using the proportion of species with a hypometabolic state compared to the assemblage. We generated maps using only the species with data from the literature review, including any species with presence or possibly presence of a hypometabolic state as presence. Then, we repeated the calculation of species richness with all the species in a grid cell to calculate the proportions of the total species in each cell with a HS.

### Distributional range analyses

In order to explore whether the species with a hypometabolic strategy had larger geographic ranges and account for phylogeny we used a phylogenetic generalized least squares (PGLS) regression. We ran five different evolutionary models (OURandom, OUFixed, BM, Pagel’s Lambda and Pagel’s Kappa) to estimate the relationship between the continuous probability estimates for aestivation, hibernation, and hypometabolism with distributional range size, specifically log1p(area), which computes log(1 + area). These models were run for each phylogeny. The best performance models, for both phylogenies, were Pagel’s Lambda and Pagel’s Kappa in both normality of the residuals (visually checked through histograms) and lowest AIC. Since Pagel’s Lambda was the model with the lowest values of AIC we chose this model to discuss in the manuscript. We used the ‘phylolm’ package (Ho & Ane, 2014). Finally, we counted the threatened category of the species in the literature dataset and discussed the implications for conservation, particularly regarding the upcoming environmental changes.

### Amphibian assemblage climatic resilience

According to our hypothesis, the greater the proportion of species with hypometabolism, the higher climatic resilience the assemblage would have. To test this idea, we quantified dissimilarities between the baseline (1990–2020) and future climate scenarios (2070–2100) by calculating the standardized local anomalies (i.e., how atypical is a value compared with the historical variability) per raster pixel (∼ 1 km^2^), using the standardized Euclidean distance (SED) (Williams *et al*., 2007). In other words, the temporal differences for each climate parameter are standardized using the local inter-annual standard deviation for that parameter. This method is able to provide information on the extent of the change in each location in absolute terms, therefore it gives direct information on the magnitude of climate change to which the assemblage might be exposed in a particular grid cell (Williams *et al*., 2007; Taheri *et al*., 2024a).

We focused on the monthly values of the multi-model mean of precipitation, and maximum and minimum of temperature from the experiments from KNMI 2023 standard set of CMIP6 General Circulation Models. Historical and future change scenarios were obtained from the Climate Explorer (https://climexp.knmi.nl). We choose a middle shared socioeconomic pathway: the regional rivalry climate change scenario ssp370 or the “rocky road”, where countries prioritize national interests over global cooperation thus leading to an increase in greenhouse gases. We use the mean of all 24 models for historical and future scenarios of climate change (CCESS-CM2, ACCESS-ESM1-5, AWI-CM-1-1-MR, BCC-CSM2-MR, CMCC-CM2-SR5, CNRM-CM6-1-HR-f2, CNRM-CM6-1-f2, CNRM-ESM2-1-f2, CanESM5-CanOE-p2, CanESM5, EC-Earth3-Veg, EC-Earth3, FGOALS-g3, GFDL-ESM4, GISS-E2-1-G-p3, INM-CM4-8, INM-CM5-0, IPSL-CM6A-LR, MIROC-ES2L-f2, MIROC6, MPI-ESM1-2-HR, MPI-ESM1-2-LR, MRI-ESM2-0, and UKESM1-0-LL-f2). In order to estimate the SED, we employed the *sed* function in the ‘climetrics’ package v.1.0-15 (Taheri *et al*., 2024b), analyses were performed using the platform R version 4.4.2 (R Core Team, 2024).

To explore where would be located the most vulnerable amphibian assemblages we first averaged climatic standardized distance per grid (1° × 1°) to match the scale of the species richness maps. We generated a heat map of the climate change magnitude through SED against the proportion of species with a HS (only literature data) under the assumption that the higher the proportion of species with a HS, the higher the expected resilience to climatic change. In order to localize the most vulnerable assemblages we selected the sites with highest increase in climate variance (> 2 of SED) and low proportion of species with hypometabolism (≤ 0.5). Afterwards, we plotted the selected assemblages on a map, we also included grid cells with the 10th highest percentile of the anomaly scores which corresponds to large changes in temperature and precipitation.

## Results

The systematic search rendered 280 articles for aestivation and 650 for hibernation. We found information about a hypometabolic strategy for 857 amphibian species (Table S1). From these species, 648 had some information about aestivation, 342 with presence, 235 with the possibility of presenting that strategy, and 52 as non-aestivating species. In the case of hibernation, we found information for 479 species, 308 with presence, 62 species with possible hibernation, and 48 that showed no hibernating strategy. The representativity of this data set is as follows: Gymnophiona, one of nine families (11%); Caudata, ten of ten (100%); and Anura, 28 of 59 (48%). Most of these species that present aestivation (389) are categorized as Least Concern (LC), 20 species are classified as Endangered (EN), 9 species from *Litoria* and *Xenopus* genera, are in the category of Critically Endangered (CR) and only *Anaxyrus baxteri* is classified as Extinct in the Wild (EW). In the case of hibernation, the proportion of species is similar with most of the species (242) categorized as LC, 19 species classified as EN, 3 as CR, two species, *Cynops wolterstorffi* and *Rheobatrachus silus*, as Extinct (EX) and *A. baxteri* as EW (Table S1; Figure S5).

### Ancestral states

The accuracy of the traits inferred for the test dataset using OC-SVM compared to data from the literature was high, with accuracies of 73% and 88 % for aestivation and 72% and 88 % for hibernation, for Jetz & Pyron and Portik *et al*. phylogenies, respectively (Figures S1–S2); in addition, all confirmed aestivation and hibernation presences based on the literature were predicted as presences by OC-SVM with > 80% probability.

The ancestral state reconstruction analyses for the 100 Jetz & Pyron (2018) trees showed a high uncertainty regarding the state for the MRCA of all amphibians, with an estimate close to 50/50 (presence/absence) for each strategy: aestivation 48.33/51.67 (SD ± 9.07), hibernation 51.51/48.49 (SD ± 14.70), and hypometabolism 48.27/51.73 (SD ± 9.83) (Figure 3; Table 1). It is worth mentioning that the standard deviation is the same for presence and absence since the values for each tree node are complementary. With the Jetz and Pyron (2018) phylogeny, the probability of the presence of a hypometabolic state for either aestivation or hypometabolism increases rapidly towards the present, being the most probable state for the MRCA of each amphibian Order: probabilities for Anura (frogs) of aestivation 62.79 (SD ± 11.23) and hypometabolism 73.09 (SD ± 13.73); for Caudata (salamanders) 68.34 (SD ± 11.97) and 78.66 (SD ± 12.30); and for Gymnophiona (caecilians) 71.72 (SD ± 16.22) and 77.26 (SD ± 14.74) (Table 1). But this was not the case for hibernation where Anura (53.97 ± 14.47) and Caudata (67.23 ± 18.49) tended for a presence, while in Gymnophiona hibernation seems more likely to be absent (64.74 ± 15.44) than present (35.26 ± 15.44). Analyses using the Portik *et al*. (2023) phylogeny also support the presence of a HS in the ancestor of Anura (70.9/29.1 ± 14.6), but provided more mixed results for hibernation (47.9/52.1 ± 9.90) and aestivation (50.2/49.8 ± 2.30), specifically. Nonetheless, the internal nodes for both phylogenies tended to be predicted to have presence of hibernation even in the superfamilies (Table 1). On average, 362 (SD ± 86) gains were inferred for aestivation and 262 (SD ± 62) for hibernation; 327 (SD ± 80) and 248 (SD ± 84) losses were inferred for aestivation and hibernation, respectively. In the combined hypometabolism, 283 (SD ± 76) gains and 250 (SD ± 69) losses were inferred. In all three strategies, there were more gains than losses (Figure S6). For the Portik *et al*. (2023) phylogeny, on average, 200 (SD ± 36.1) gains were inferred for aestivation and 163 (SD ± 34.2) for hibernation; 150 (SD ± 41.7) and 170 (SD ± 35.4) losses were inferred for aestivation and hibernation, respectively. For hypometabolism, 161 (SD ± 45.1) gains and 151 (SD ± 32.7) losses were inferred (Figure S7).

**Figure 3.**
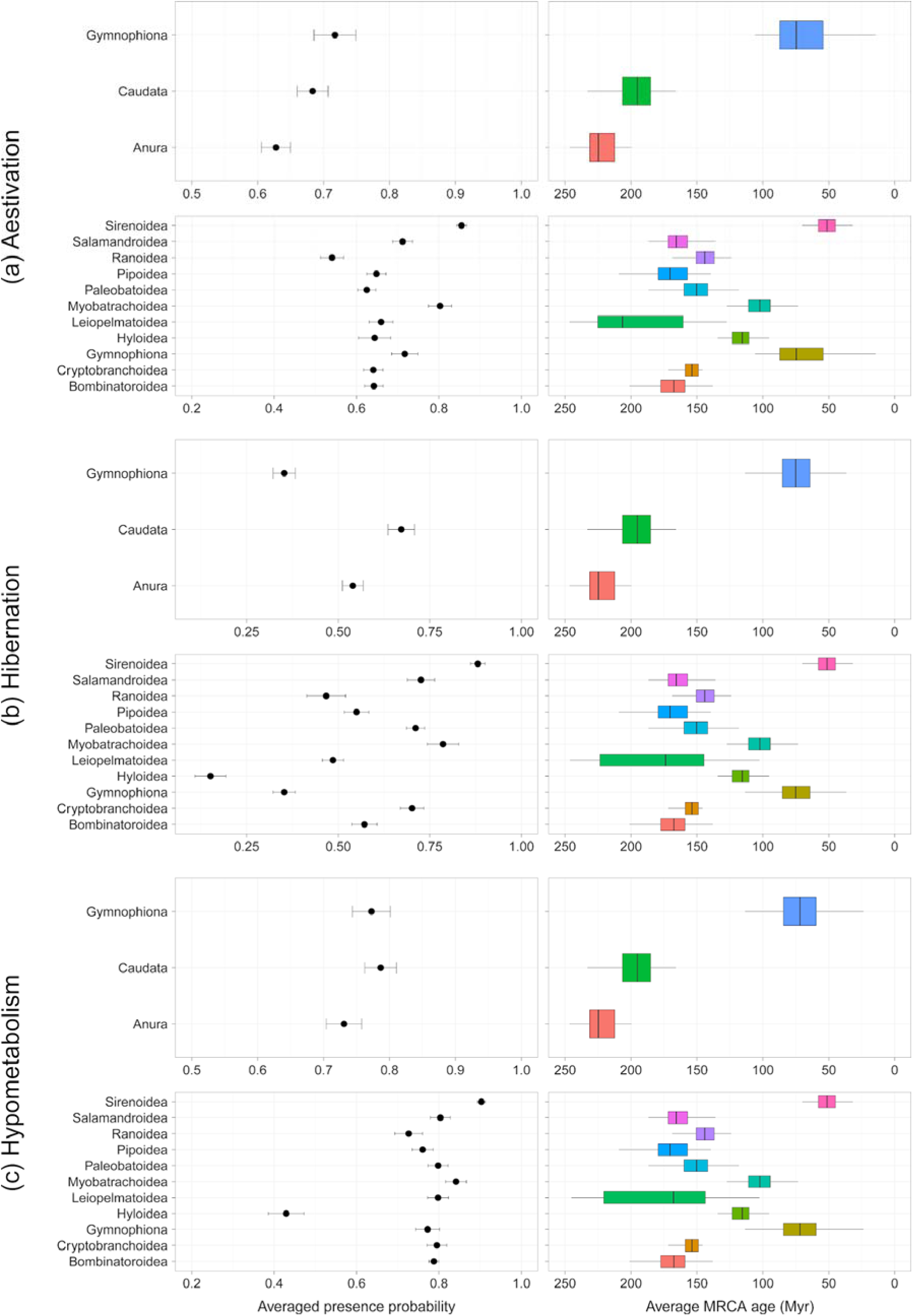
Probabilities from the ancestral state reconstruction and age node estimation with the selected from Jetz and Pyron (2018) phylogeny for A) aestivation, B) hibernation and C) hypometabolism. Result from the MCMC approach to sample character histories from their posterior probability distribution (multiSIMMAP analyses) for 100 trees.

**Table 1.**
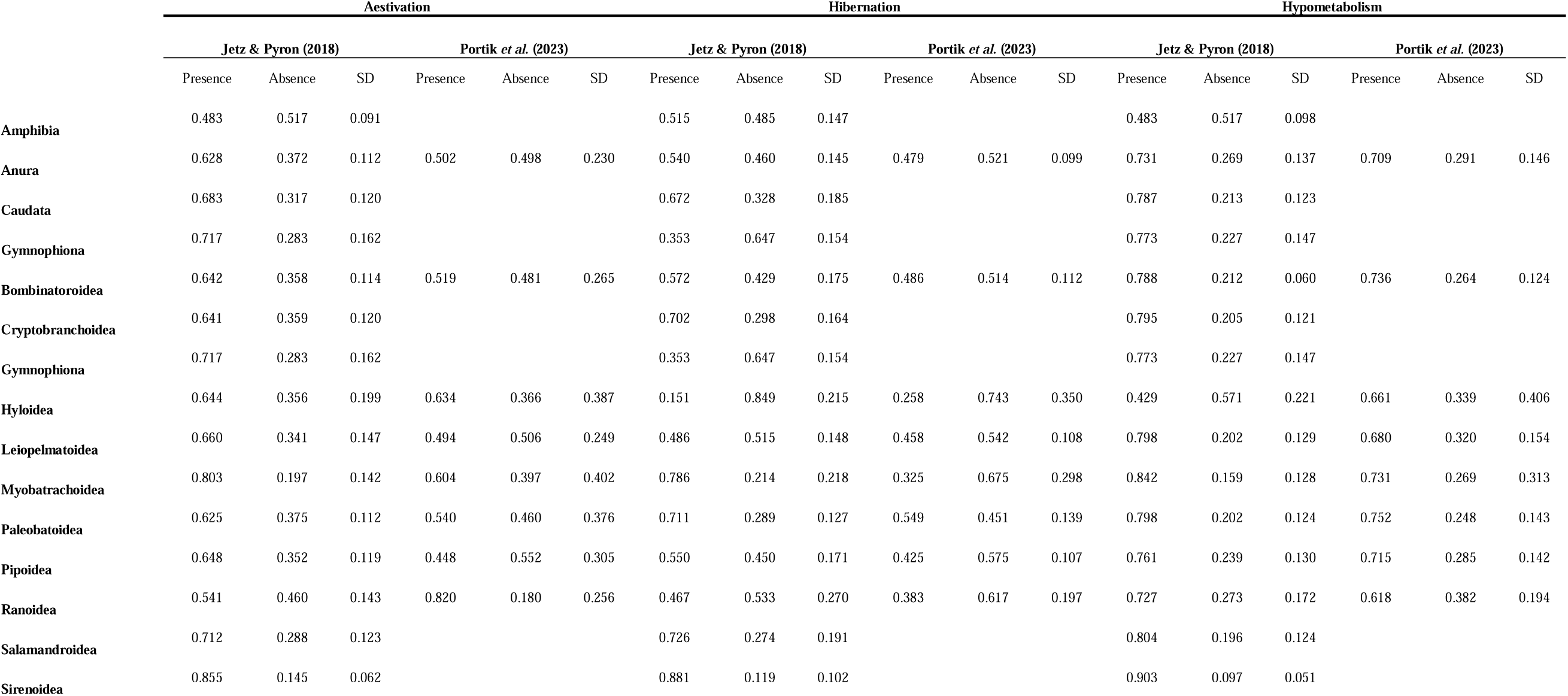
Comparative table from the ancestral state reconstruction for aestivation, hibernation and hypometabolism for all amphibians with Jetz and Pyron (2018) phylogeny and for just Anura with Portik *et al*. (2023) phylogeny.

For aestivation it is possible to observe a complex pattern of gains and losses of the trait (Figure 4), where most of the clades have representative lineages of both, although there are a few groups exhibiting only presence such as the Pelodryadinae subfamily (i.e., *Cyclorana*), and some genera within the subfamilies Hyloxalinae and Mantellinae, and the Ceratophryidae family. The transitional evolution of aestivation is likely to be complex, for example being the most likely ancestral state in groups like Myobatrachoidea (> 80% with Jetz & Pyron phylogeny and > 60% with Portik *et al*. phylogeny), but for Ranoidea although probabilities are above 50 and 80% there seems to be multiple independent losses (Figure 4). For hibernation there are only a few groups that show a high probability for the presence of hibernation as an ancestral state, such as Caudata, as well as in some anurans, including Myobatrachoidea (∼ 80% and 78%), Paleobatoidea (∼ 50% and 80%) and Pipoidea (∼ 64% and 55%), while Hylodea 15% and 25% seems to have a very low probability of presence of hibernation (Table 1, Figures 4 and S8). Meanwhile groups such as Sirenoidea, Salamandroidea, Myobatrachoidea, and Bombinatoroidea showed remarkably high probabilities of presence of hypometabolism (Table 1). In the case of the Portik *et al*. phylogeny all anuran superfamilies showed > 60% probability of hypometabolism.

**Figure 4.**
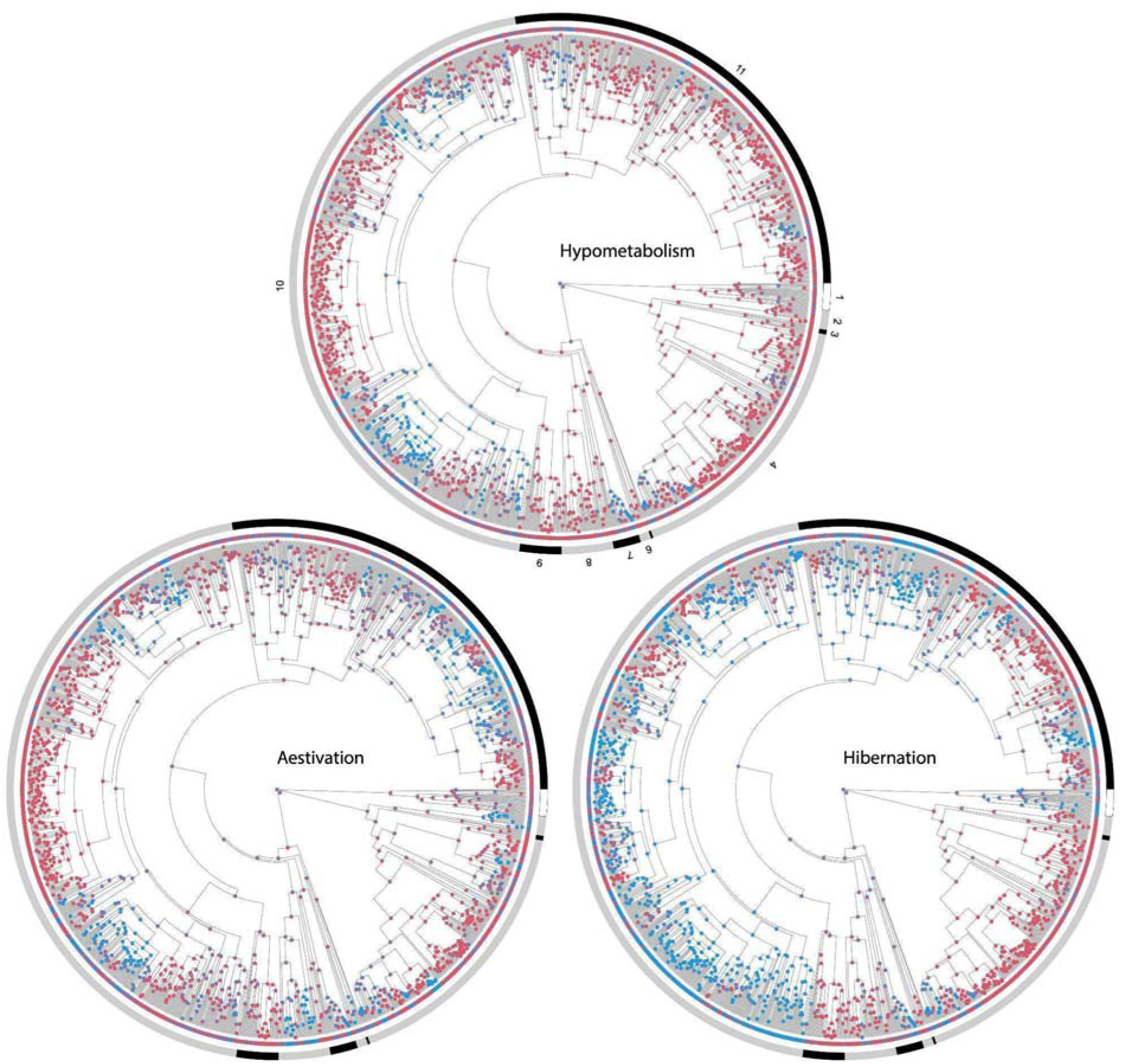
Stochastic character mapping for the hypometabolic strategies in amphibians (red = presence; blue absence) based on a single tree. The internal colored ring depicts the current state of each tip, while the external ring indicates major groups as featured in Jetz and Pyron (2018): 1. Gymnophiona, 2. Cryptobranchoidea, 3. Sirenoidea, 4. Salamandroidea, 5. Leiopelmatoidea, 6. Bombinatoroidea, 7. Pipoidea, 8. Paleobatoidea, 9. Myobatrachoidea, 10. Hyloidea, and 11. Ranoidea.

At a broad scale, based on the selected tree only for the Jetz & Pyron phylogeny, the proportion of ancestral nodes with aestivation decreased during the Jurassic when the Earth’s surface temperature was transitioning to a cool-house (ca. 175–160 Myr), to posteriorly recover again in the transition to a warm-house (ca. 160–150 Myr). Unexpectedly, aestivation’s greatest depression was inferred during a warmer period between 150–125 Myr, with a posterior recovery and fluctuating but steady trend to the present. On the other hand, the hibernation trend was stable during the cooled period in the Jurassic, with a drop during the transitional period to warmer temperatures 160 Myr, with a posterior recovery but reduction again during the remaining Mesozoic, and a recovery during the transition to a cooler Earth in the past 50 Myr (Figure 5).

**Figure 5.**
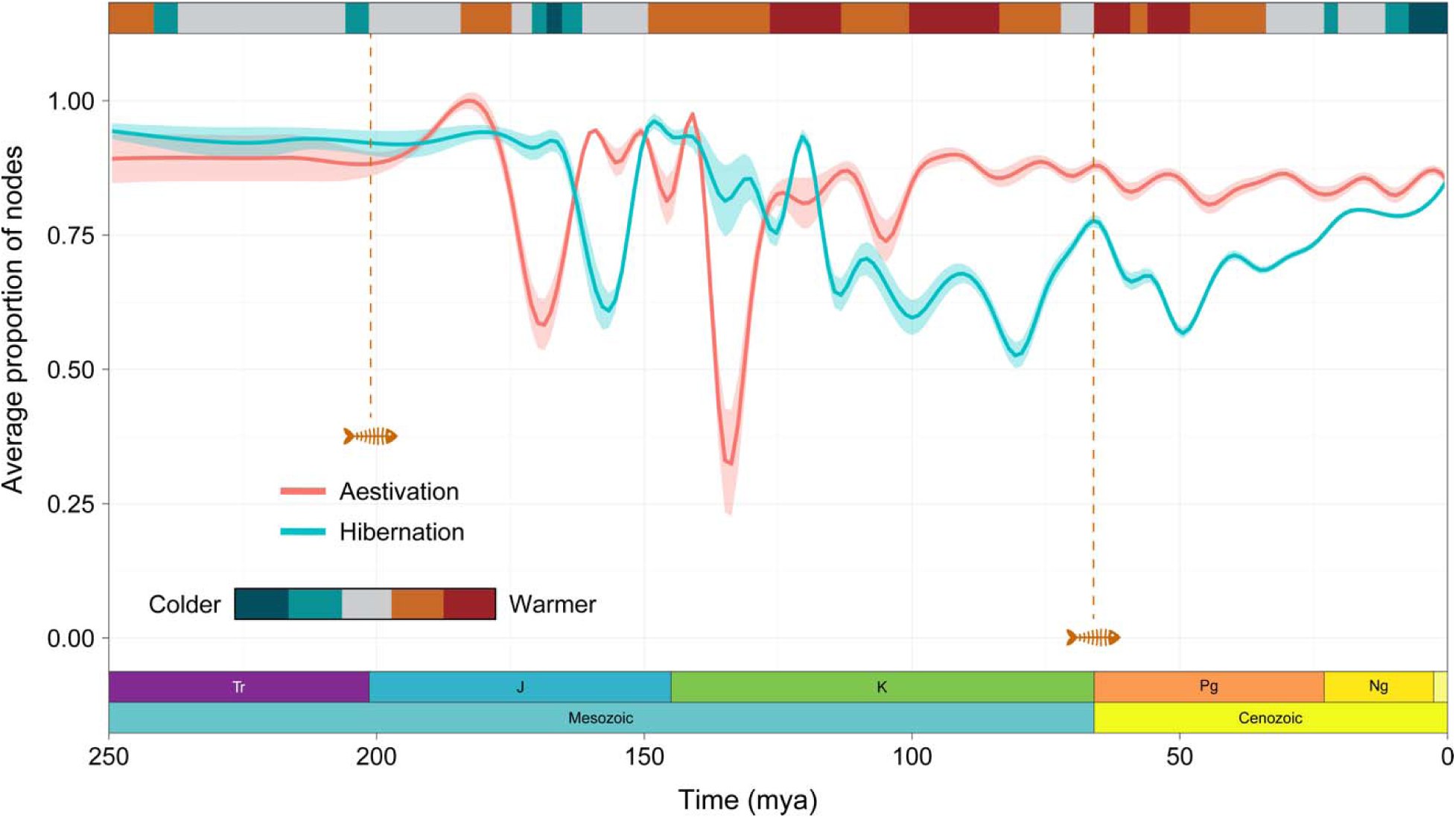
Average proportion of nodes that were inferred as presence as the most likely state for one of the two hypometabolic strategies, according to the joint ancestral state reconstruction for selected tree for the Jetz & Pyron (2018) phylogeny. The temperature bar is the same as featured in Judd *et al*. (2024). Curves were smoothed using the smoothing splines method and shaded areas are derived from the confidence interval (95%). Dotted lines with the fish skeleton indicate major extinction events, the Triassic–Jurassic (Tr-J) 200 Myr, and the Cretaceous–Paleogene (K–Pg) 66 Myr.

For Portik *et al*. phylogeny and based on the selected tree only, a similar pattern as with Jetz & Pyron phylogeny occurred during the Jurassic but with the interesting fact that the average of presence decreases to zero, to posteriorly recover again in the transition to a warm-house (ca. 150 Myr). Another aestivation depression was inferred during a warmer period between ∼ 95–85 Myr, with a posterior recovery, and another depression ∼ 70 Myr and finally a recovery fluctuating but steady trend to the present. While aestivation was in depression during the Jurassic the proportion of ancestral nodes with hibernation increased. The hibernation trend showed gains and losses (but never as deep as in aestivation) during the remaining Mesozoic, and a stable trend during the transition to a cooler Earth in the past 50 Myr (Figure S9).

### Geographic distribution of species with hypometabolism

As mentioned in the methods, here we are only using literature data. The greatest number of aestivating species are located in the Eastern United States, Northern and Eastern Australia, parts of Indonesia, the Dry Diagonal in South America, and in South Africa and Zimbabwe (Figure S10). In terms of the proportion of species in a community with aestivation, the pattern is much different, highlighting regions of the world with arid climates such as southwestern US, Botswana, parts of the Middle East and northern Africa, New Zealand and much of Australia (Figure S11). Hibernating species richness is predicted to be highest in the eastern United States, parts of China and Europe (Figure S12), yet communities with the highest proportion of species that hibernate clearly predominate in northern latitudes and in the Himalayas (Figure S13).

### Distributional range analyses

The results of the phylogenetic generalized least squares (PGLS) regressions based on the Lambda model showed no relation between the probability of any hypometabolic state and distribution range size on both phylogenies. This can be observed in the really low values of the adjusted R^2^ (Table 2), meaning that the unexplained variance and error of the model is high.

**Table 2.** Summary of the phylogenetic generalized least squares (PGLS) regressions based on the Lambda model. Colums represent the average and standard deviation of each resulting parameter.

| | $\sigma^2$ | Lambda | logLik | Adjusted R<sup>2</sup> |
| --- | --- | --- | --- | --- |
| **Jetz & Pyron (2018)** |  |  |  |  |
| Aestivation | 0.070 ± 0.004 | 0.670 ± 0.016 | -15299.414 ± 10.069 | 0.071 ± 0.001 |
| Hibernation | 0.064 ± 0.004 | 0.644 ± 0.016 | -15181.902 ± 8.571 | 0.106 ± 0.001 |
| Hypometabolism | 0.066 ± 0.004 | 0.652 ± 0.016 | -15254.762 ± 9.189 | 0.084 ± 0.001 |
| **Portik *et al.* (2023)** |  |  |  |  |
| Aestivation | 0.118 ± 0.008 | 0.856 ± 0.010 | -9431.548 ± 4.733 | 0.115 ± 0.001 |
| Hibernation | 0.122 ± 0.008 | 0.842 ± 0.011 | -9603.590 ± 4.116 | 0.034 ± 0.001 |
| Hypometabolism | 0.103 ± 0.007 | 0.829 ± 0.012 | -9380.125 ± 3.979 | 0.139 ± 0.001 |

### Amphibian assemblage climatic resilience

The results of the standardized climate anomalies showed higher magnitudes of climate change occur in the tropics (Figure 6). However, most sites will experience low climate change; these sites have a wide range of proportions of species with a HS (Figure 6a). We consider that most vulnerable assemblages are sites with high climate variance (> 2 of SED). If we take the subset of these that contain a low proportion of species with hypometabolism (≤ 0.5), there are a total of 238 assemblages, all of which are located in the tropics (Figure 6b). A few of these sites do have a high proportion of species with a hypometabolic state, and, therefore, a climatically resilient assemblage.

**Figure 6.**
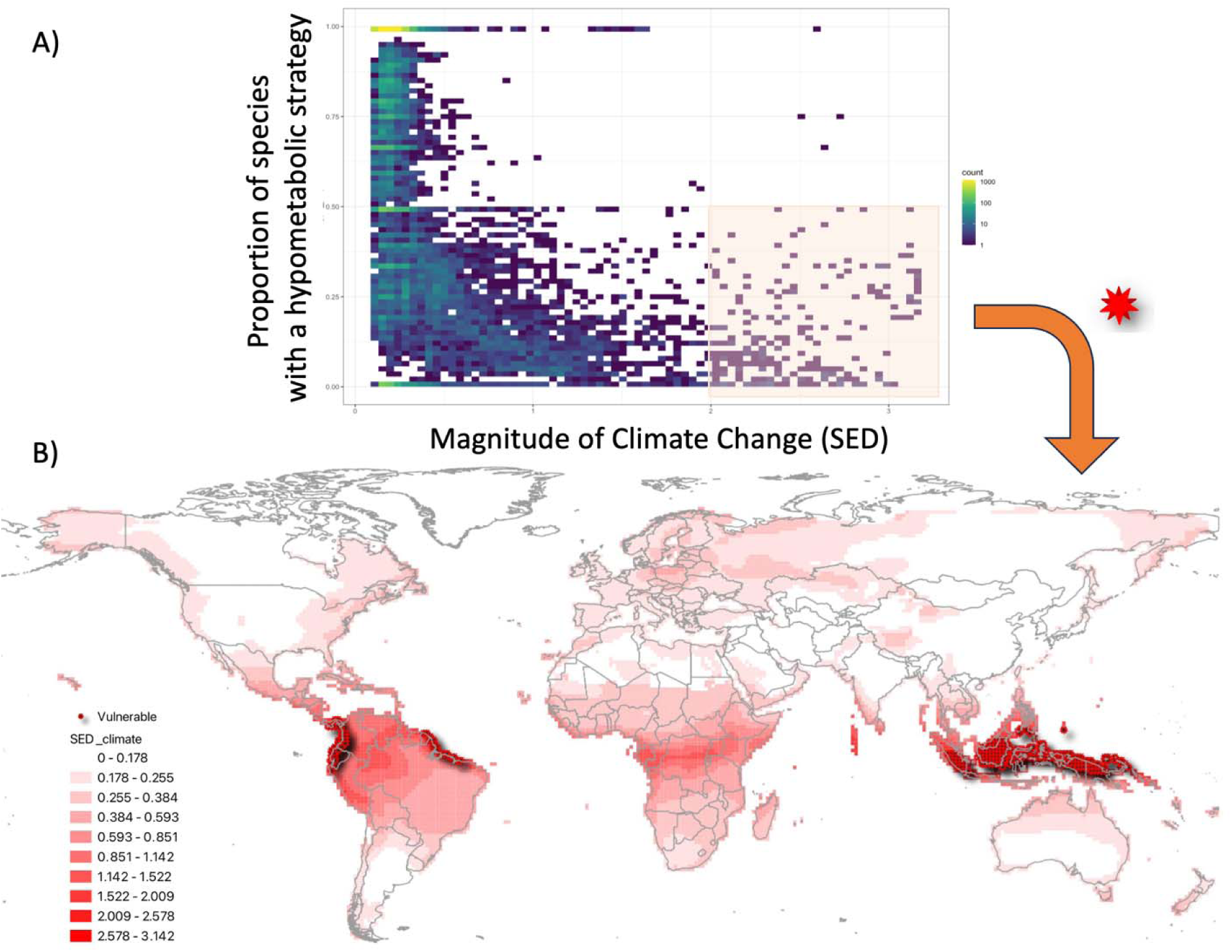
Standardized Euclidean distance based on temporal differences for each climate parameter which are standardized using the local inter-annual standard deviation for that parameter in both periods of time, in this case t1 (1990-2020) and t2 (2070-2100), this can be interpreted as magnitude of climate change. The higher the standardized local anomaly scores the larger the predicted changes in temperature and precipitation. The most vulnerable amphibian assemblages based on the low proportion of species with a HS, only literature data (bottom right in A) are highlighted with a red star in B.

## Discussion

With more than 8,800 described amphibian species, we only recovered information for fewer than 10%, clearly indicating that more information about the ecology of amphibian species is needed. This is troublesome to say the least, not only because it increases the uncertainty in the analyses (see below) but also because it means that for the majority of the species, we do not have the minimum information to establish effective conservation plans and we will need to rely on assumptions derived from < 10% of each group or class. Nevertheless, we note that these 857 species include a variety of distinct lineages from caecilians to salamandrids, pipids, and hylids to name just a few. Another aspect to take into account is the inherent geographical bias of the data, where most of the data is from the USA, Australia, New Zealand, and the European region. Although, it is worth mentioning that other regions also stand out, such as South America and South Africa, probably due to the extreme seasonality and the number of species that are known to estivate in these regions.

### Reconstruction of ancestral states

At the root of the tree for all amphibians the ancestral state reconstruction showed high uncertainty for the presence of any hypometabolic state (∼ 50%). However, at the order and superfamily levels, the ancestral state reconstruction analyses provide support for the presence of hypometabolism, > 70% for all three orders in Jetz & Pyron and in Anura for Portik *et al*. phylogenies. Support was low for the presence of hibernation as an ancestral state in Gymnophiona (35%) and moderate to low for the other two orders, 67% in Caudata, and 54% and 49% for Anura in the Jetz & Pyron and Portik *et al*. phylogenies, respectively. Support for the presence of aestivation was moderate to high in all three orders, ranging from 63% and 50% for Anura (on each phylogeny, respectively), 68% for Caudata, and 71% for Gymnophiona.

Based on the probability of presence at the Order level, the HS seem to have been secondarily lost multiple times within the amphibian tree; the complex pattern presented in the different reconstruction analyses prevents us from drawing a general inference for the whole group. As we acknowledged, there is very limited data regarding the absence of either aestivation or hibernation, so additional data and analyses will be necessary to confidently assign losses of the trait. For example, our OC-SVM model yielded several “false positive” species. In this regard, if the climate is indeed highly influencing the presence of a hypometabolic strategy, and it is somehow predictable, those “false positive” could be “true positive” reflecting a bias in the lack of knowledge about the species’ ecological traits (Hortal *et al*., 2015).

There are differences in the reconstruction of ancestral states between aestivation and hibernation, and these are likely a reflection of their different evolutionary histories, even though aestivation and hibernation are both cases of an adaptive dormancy (Navas & Carvalho, 2010), the mechanisms that trigger each of the different states of hypometabolism are completely different. On the one hand, aestivation seems to be triggered by lack of water and excessive heat, on the other, hibernation seems to be triggered by low temperatures, and it is commonly associated with accessible water. Nonetheless, it is possible that the regulatory mechanisms responsible for depressing metabolisms are the same between aestivation and hibernation (e.g., methylation of DNA) (Hawkins & Storey, 2020; Bell & Hellmann, 2025) but the triggering event is different in each case. So, our results seem to agree partially with the proposal of Storey & Storey (1990) in that if having an hypometabolic state is an ancestral state, therefore the mechanisms and regulatory signals that are activated during metabolic depression have a common molecular basis for all states of metabolic depressions, including aestivation, hibernation, and some types of anoxybiosis, probably even cryobiosis. It might be possible to go even further and predict that torpor also relies on the same mechanisms; however, little is known in this aspect for amphibians since most of the literature about torpor is actually referring to winter torpor, here classified as hibernation (Boutilier *et al*., 1997; Wilkinson *et al*., 2017) but see Castanho and De Luca (2001).

### Distributional range analyses

The results of the phylogenetic generalized least squares (PGLS) regressions showed no relation between the probability of having a HS and the distribution range size. We expected that species with a hypometabolic strategy would have significantly larger distribution ranges because having a hypometabolic strategy allows individuals and populations to overcome harsh environments since they do not have to ‘suffer’ those conditions. Therefore, this ability could allow them to disperse to stressful or unfavorable environments because they are not limited by narrow environmental constraints. Nevertheless, when accounting for phylogeny no relationship was shown.

### Geographic distribution of species with hypometabolism

The pattern of aestivating species richness (Figure S9) is really striking in some parts of the world such as in the east coast of the United States where there is a high number of species that present aestivation. In other regions such as Australia, South Africa, the arid zones of North America and the Arid Diagonal in South America, the high number of species that present aestivation is unsurprising due the high precipitation seasonality in those regions. The pattern of the proportion of species that present aestivation (Figure S10) follows very closely the arid zones of the world (Zomer et al., 2022), with obvious exceptions where no amphibians are found such as the Sahara, Arabian and Gobi Deserts which are deemed as hyper-arid zones (Yang & Williams, 2015).

The pattern of hibernating species richness is also interesting in some parts of the world such as in the east coast of the United States and southeast China, where there is a high number of species that present hibernation (Figure S12). In other parts such as the rest of North America, Europe, and northern Asia, the high number of species that present hibernation is expected due the high seasonality in temperature in those regions as they coincide with Köppen cold climates map (Beck et al., 2023).

### Amphibian assemblage climatic resilience

It is interesting that when accounting for the variability along the periods tested (i.e., using the SED method), the results showed that the largest differences between current and future climate is going to occur in the tropics, in other words where the most atypical values will occur. These sites are going to experience the greatest change in climate, according to the models, and therefore amphibian populations living there are going to experience a substantial pressure to adapt, possibly using hypometabolic states to help them to survive. It does make sense because much of the tropics have little variability in temperature and precipitation compared to the higher latitudes where the inter-annual variation is huge. In addition, most of the species with a hypometabolic strategy currently occur outside the tropics (Figures. S10-S13). Consequently, according to our definition, the most vulnerable assemblages are located in the tropics, specifically in the lowlands. For example, soil temperature variation in tropical mountains like the Andes may reach almost 50 °C in one day (Navas, 1996), and we expect that these species are likely to demonstrate some form of hypometabolism. For example, *Atelopus carrikeri* lives at 4,500 m a.s.l. where temperatures can be high during the day but it can be trapped occasionally in ice sheets and survives (Navas, 1997). It is important to stress that high-elevation Neotropical amphibians do not belong to a single taxonomic group. Species from several divergent families such as Bufonidae, Leptodactylidae, Dendrobatidae, and Hylidae are represented at elevations over 3,000 m a.s.l. in the tropical Andes, and these high-elevation species are apparently derived from low elevation stocks (e.g., Fouquet *et al*., 2013; Mendoza *et al*., 2015). Thus, given the results of the ancestral state reconstruction, it might be possible to assume that these species were able to persist in those conditions because they have converged on the mechanisms to survive harsh conditions.

Another aspect that is worth highlighting is the temperature spectrum of activity. Navas (1996) showed that high-elevation anurans are capable of performing activities across a wide range of temperatures, however low-elevation species can be active in narrower ranges of temperature. This supports our results of assemblage climatic resilience in the sense that most vulnerable assemblages were found in tropical lowlands. Plastic responses to past environments shape adaptation to novel selection pressures (Coates *et al*., 2025), therefore it is logical to assume that if there is geographical variation in the response to climate across population, that is, not necessarily all populations aestivate or hibernate, this plasticity may give said species a chance to adapt to novel climate. It is important to stress that given the scarcity of the data it might result in a less worrisome scenario for the tropical assemblages of amphibians given the probability that more species could aestivate.

### Limitations of the analyses

Even though we have tried to account for missing data, it is important to stress that the results presented here have to be taken with caution as the information used in the analyses represents fewer than 10% of the total amphibian species described (Frost, 2024; AmphibiaWeb, 2024). Two aspects emerge from this lack of data, the first is the high uncertainty that was generated by inferring the probabilities of a hypometabolic strategy based only on climate. Although the inferences did correctly predict all of the species known to have hypometabolic strategies, such uncertainty may have contributed to noise in the dataset. The second aspect regarding uncertainty in our analyses is the assumption that species not observed to have a HS in the wild cannot present a HS. For example, *Rhinella marina* does not normally aestivate under natural conditions but when subjected to experimental conditions of water stress and temperature it does aestivate (Boutilier *et al*., 1979), but see Hilje & Arévalo-Huezo (2012). Solving this dilemma will be crucial in the years to come when planning effective conservation strategies, either by collecting more information in the field or by performing experiments. It would be ideal to have information for all genera.

### Future research directions

Aside from a call for more data on hypometabolism in amphibians, we propose a few research themes that would be fruitful to pursue in the future. First, how long can species aestivate or hibernate? A few accounts suggest that amphibians can survive five or more years while estivating (Van Beurden, 1980) and estimations suggest even more (Sadowski-Fugitt *et al*., 2012), which conflicts with common narratives that amphibians are sensitive and fragile organisms. Persisting in a hypometabolic state for years would provide impressive resiliency to sudden or sustained environmental change and may start to explain how amphibians have survived several mass extinction events. Aestivation may also help amphibians survive wildfires. Second, could resorting to aestivation or hibernation affect longevity in populations? Third, do amphibians have different levels of hypometabolism, for example, very shallow (days), shallow (weeks) or deep (months to years)? In this sense should we classify torpor as an actual hypometabolic state? Are the molecular mechanisms the same? Fourth, what are the physiological mechanisms of the species that live in mountains, such as the Andes, where they can experience an oscillation of temperature of more that 40 °C in one day? Finally, in the era of huge environmental changes, will the limits of hypometabolic ability be reached in the near future (Macip-Ríos *et al*., 2023)? Would having a hypometabolic strategy give populations resilience to face coming changes in climate?

Clearly the major drawback of the analyses and results presented here are the number of species used for the analyses. Therefore, we call on our colleagues to report whether or not a species goes into a hypometabolic state, and if possible, to test them. A review of amphibian phenology may also identify additional cases of aestivation or hibernation.

## Supplementary material

**File S1. Additional methods and results.** This file contains additional results.

**File S2. R scripts used for the analyses performed.**

**Table S1. Dataset of species with hypometabolic strategy information.** Compiled dataset containing the species found in literature regarding hypometabolic strategy (HS): aestivation or hibernation. HS was classified as **present** = confirmed hypometabolic state, **possible** = authors mention the possibility but it was not confirmed, **absent** = the reference clearly mentions that the species/population does not present the hypometabolic state**, unknown** = when the authors clearly mention that it is unknown. The table also lists IUCN category (2024), name in the trees (Jetz & Pyron, 2018 and Portik *et al*., 2023), and the calculated probabilities for the states.

