## Supplementary Material for "Slowing down: A macroevolutionary approach to the hypometabolic strategies of amphibians"

**Slowing down: a macroevolutionary approach of hypometabolic strategies of amphibians Supplementary material**

**Additional methods and results**

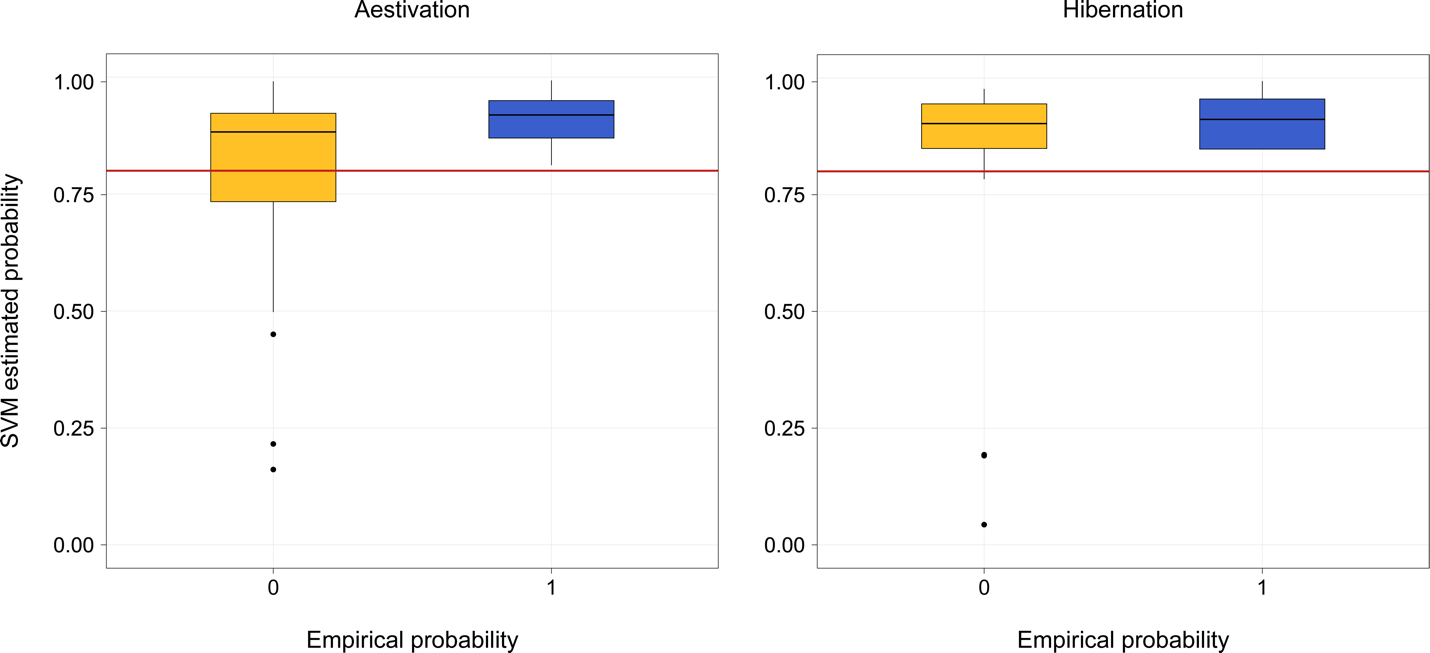

**Figure S1.** Validation of the One-Class Support Vector Machine (OC-SVM) model using a test dataset for the Jetz and Pyron (2018) phylogeny. Left: Aestivation, and right: Hibernation.

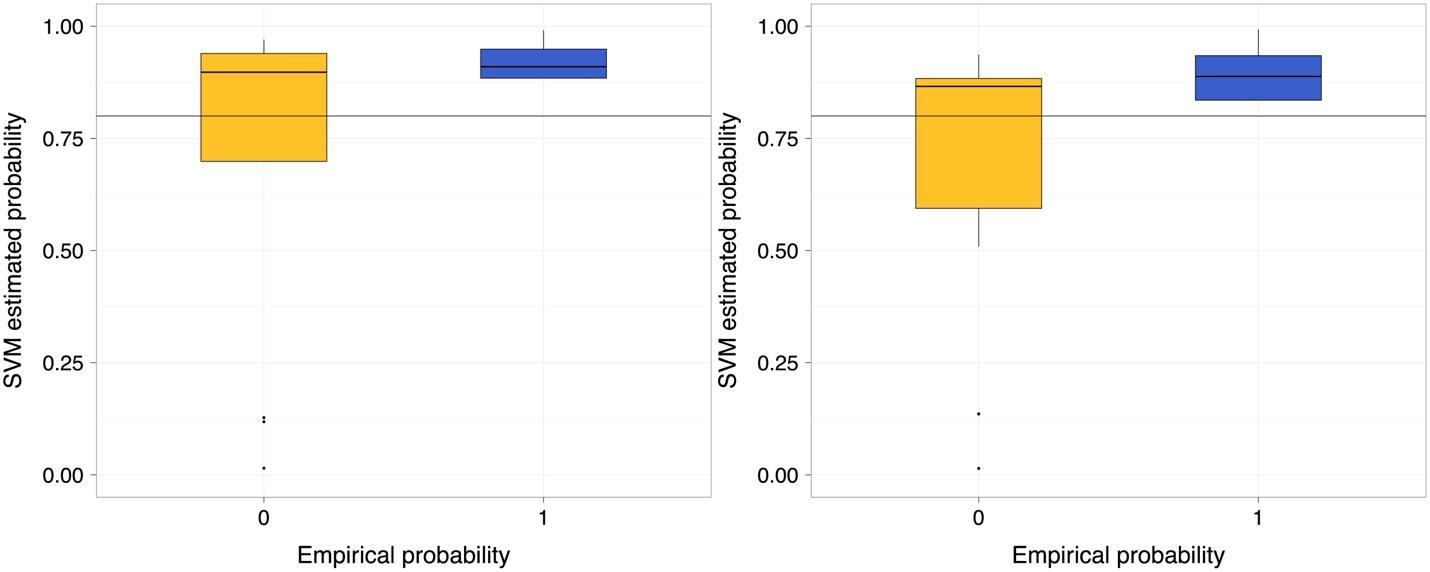

**Figure S2.** Validation of the One-Class Support Vector Machine (OC-SVM) model using a test dataset the Portik et al. (2023) phylogeny. Left: Aestivation, and right: Hibernation.

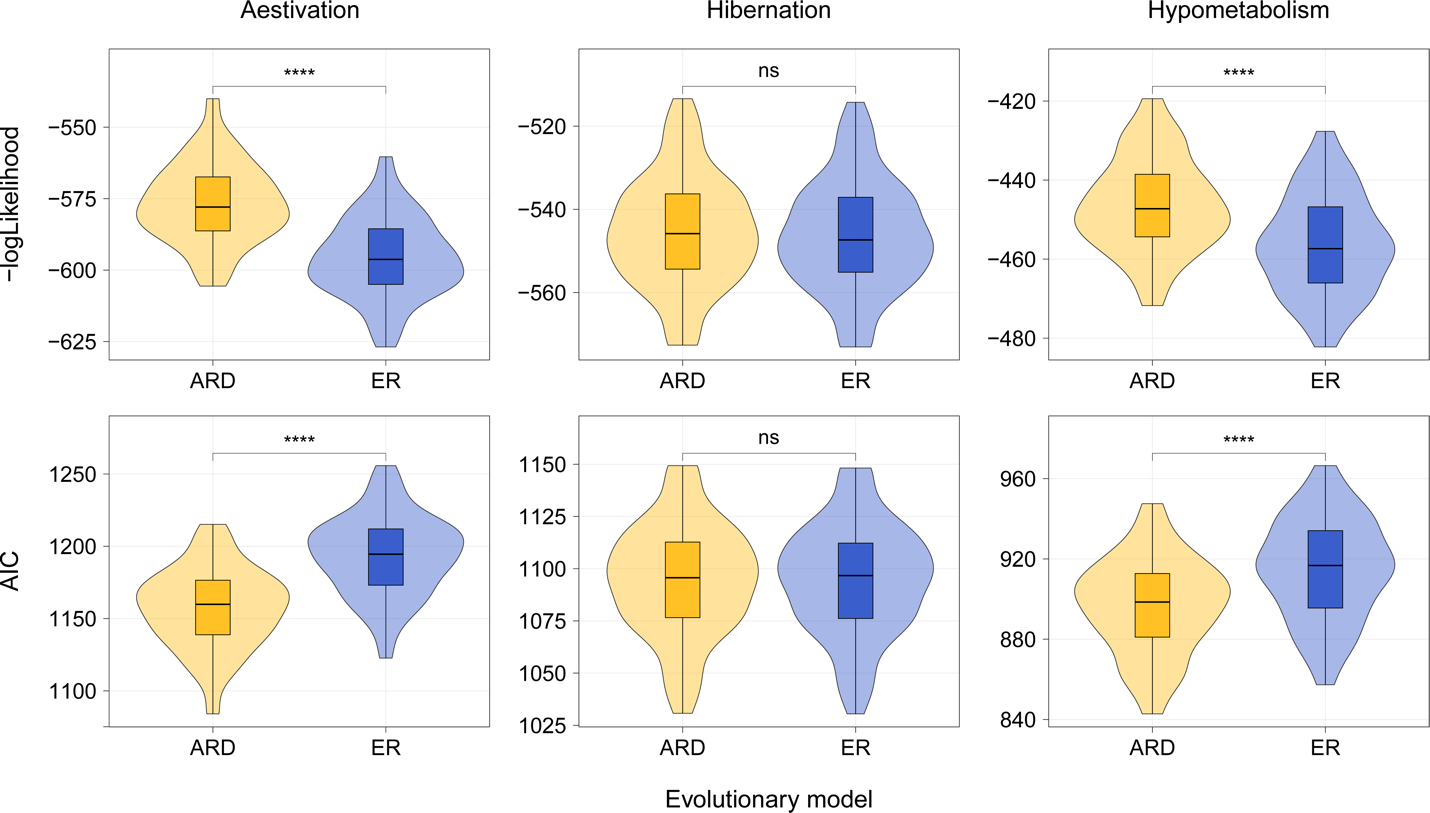

**Figure S3.** Transition models performance for the Mk model for discrete character evolution for the Jetz and Pyron (2018) phylogeny. Differences between groups were calculated using the Student’s t-test. Significance codes for p-value are as follows (ns: > 0.05 / * ≤ 0.05 / ** ≤ 0.01 / ***: ≤ 0.001 / ****: ≤ 0.0001*).

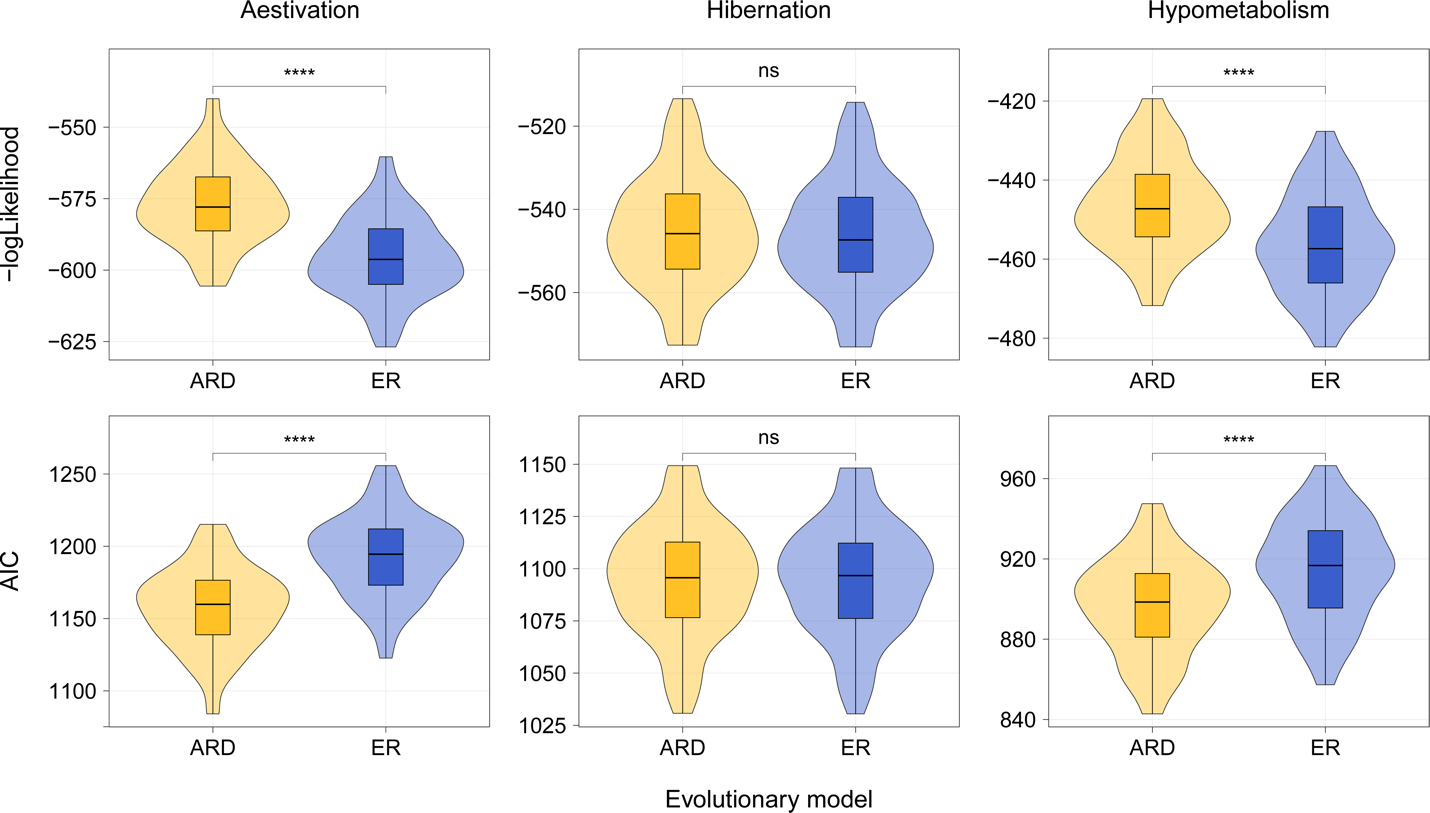

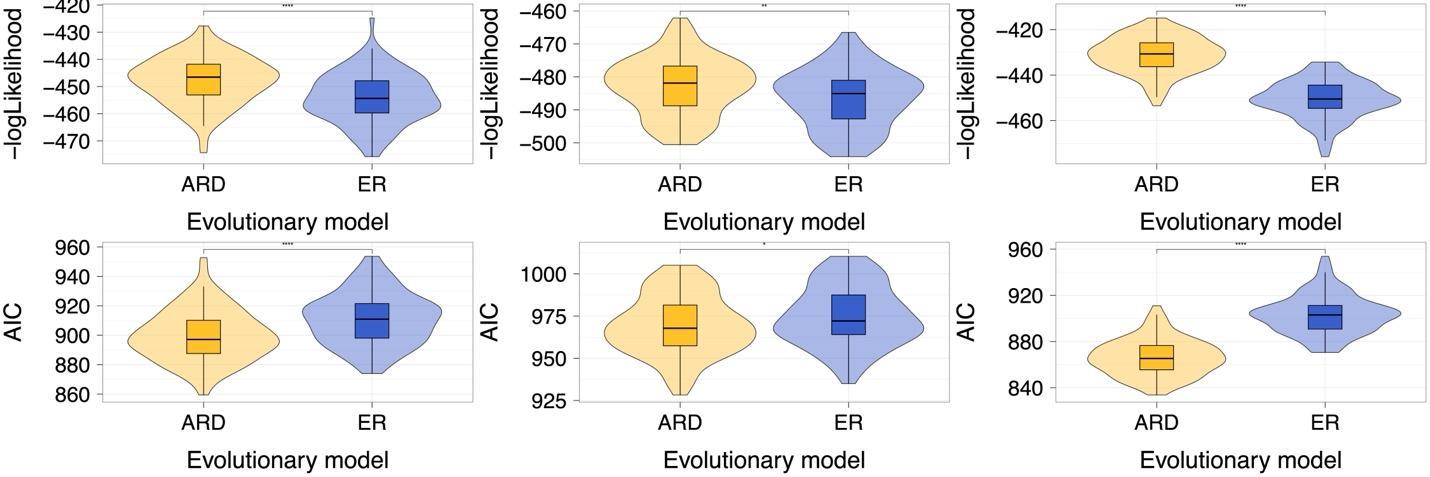

**Figure S4.** Transition models performance for the Mk model for discrete character evolution for the Portik et al. (2023) phylogeny. Differences between groups were calculated using the Student’s t-test. Significance codes for p-value are as follows (ns: > 0.05 / * ≤ 0.05 / ** ≤ 0.01 / ***: ≤ 0.001 / ****: ≤ 0.0001*).

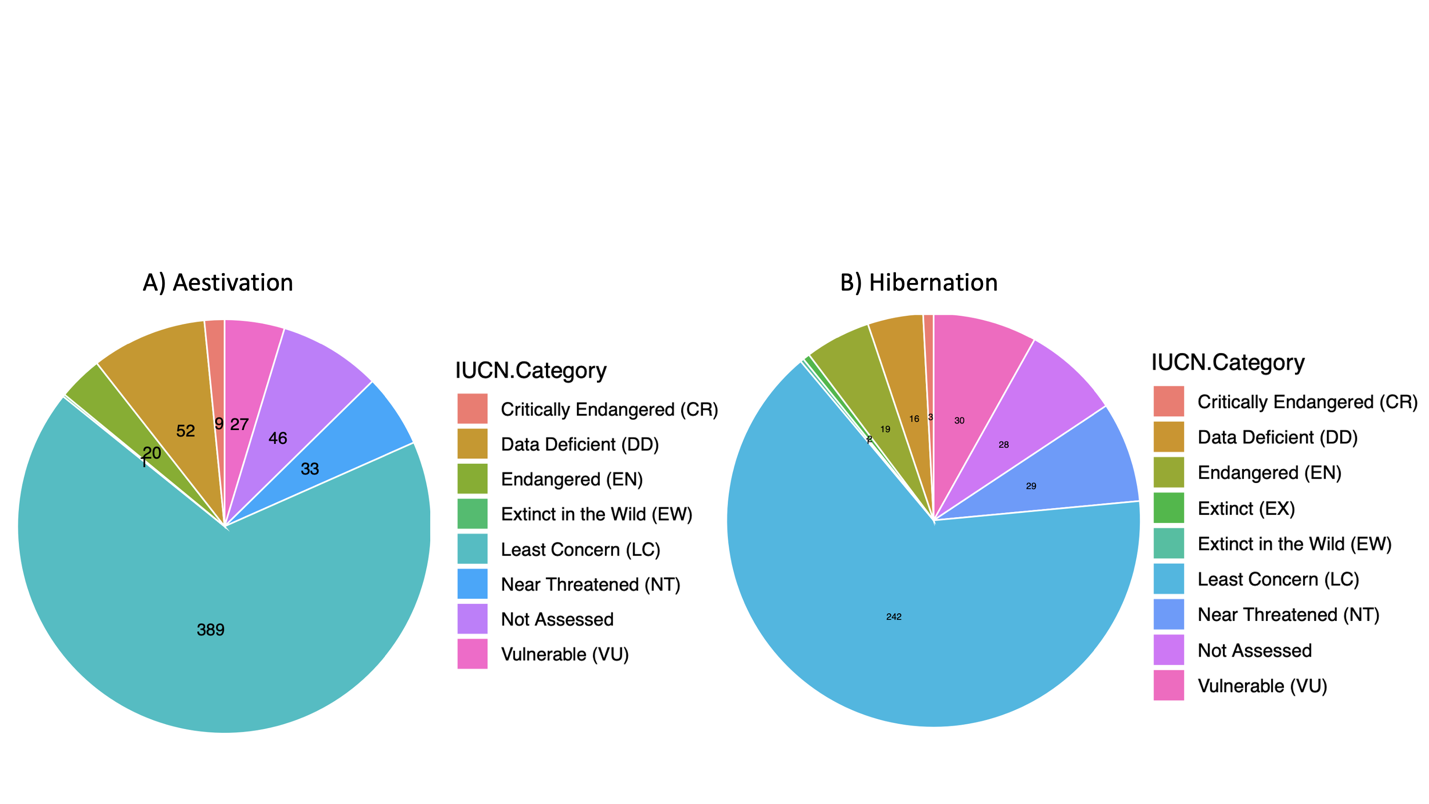

**Figure S5.** IUCN categories (IUCN, 2023) of the amphibians that presented a hypometabolic strategy. A) Aestivation, and B) Hibernation.

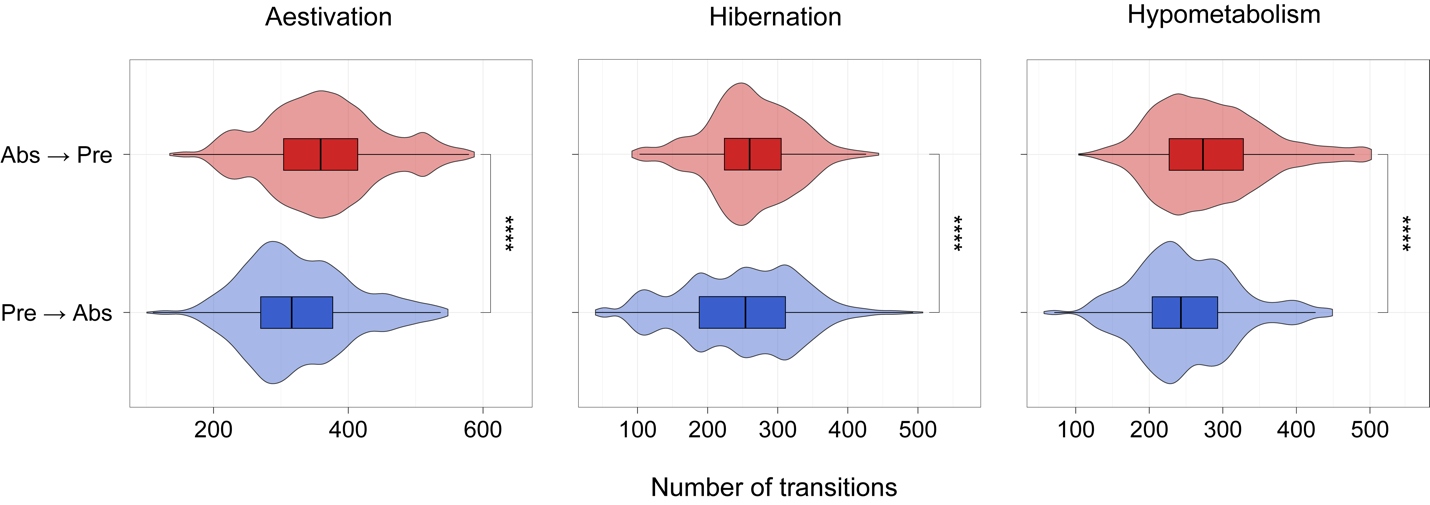

**Figure S6.** Frequency distribution for the inferred number of transitions for all the hypometabolic states for the Jetz and Pyron (2018) phylogeny. Differences between groups were calculated using the Wilcoxon test. Significance codes for p-value are as follows (ns: > 0.05 / * ≤ 0.05 / ** ≤ 0.01 / ***: ≤ 0.001 / ****: ≤ 0.0001*).

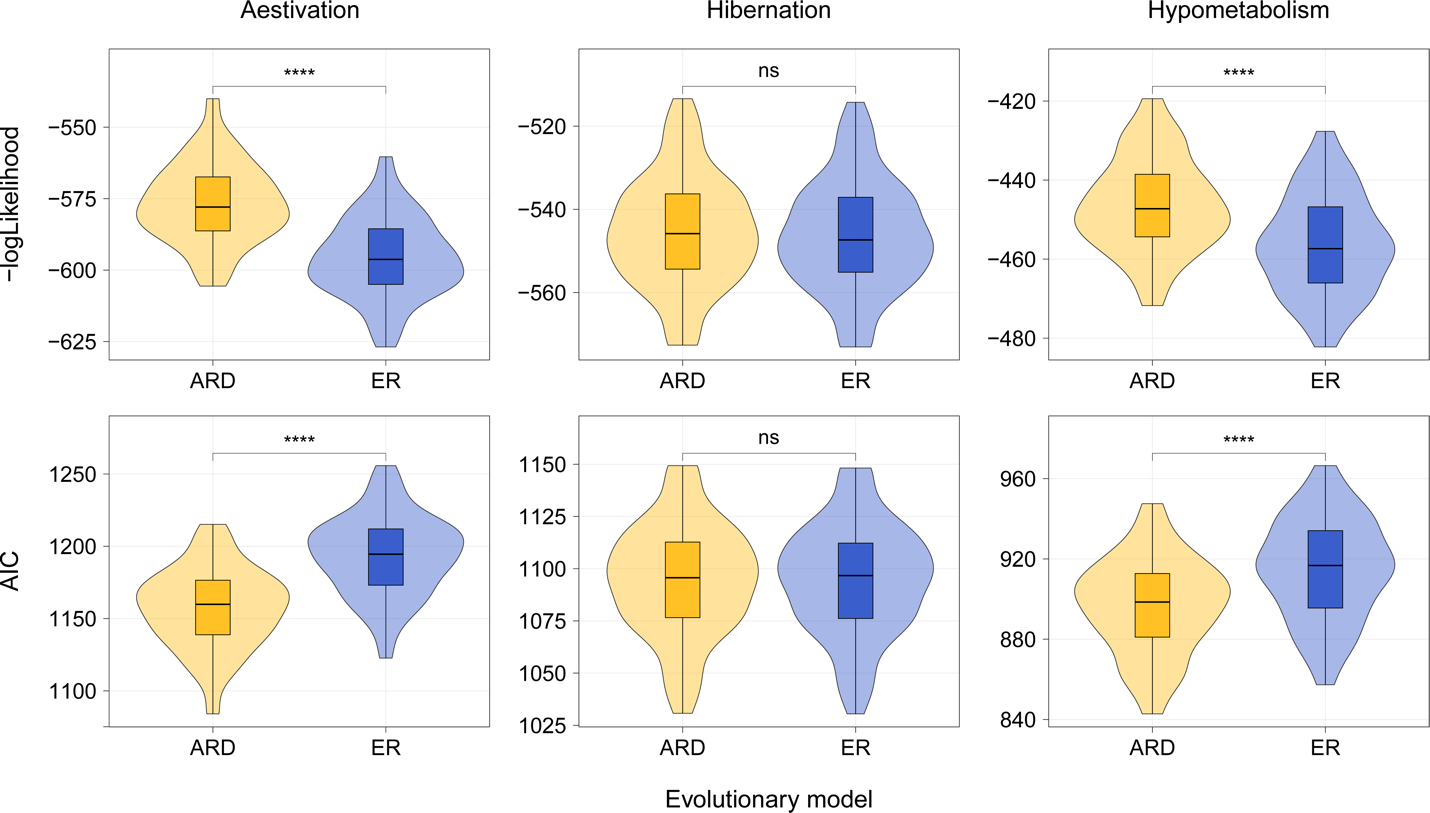

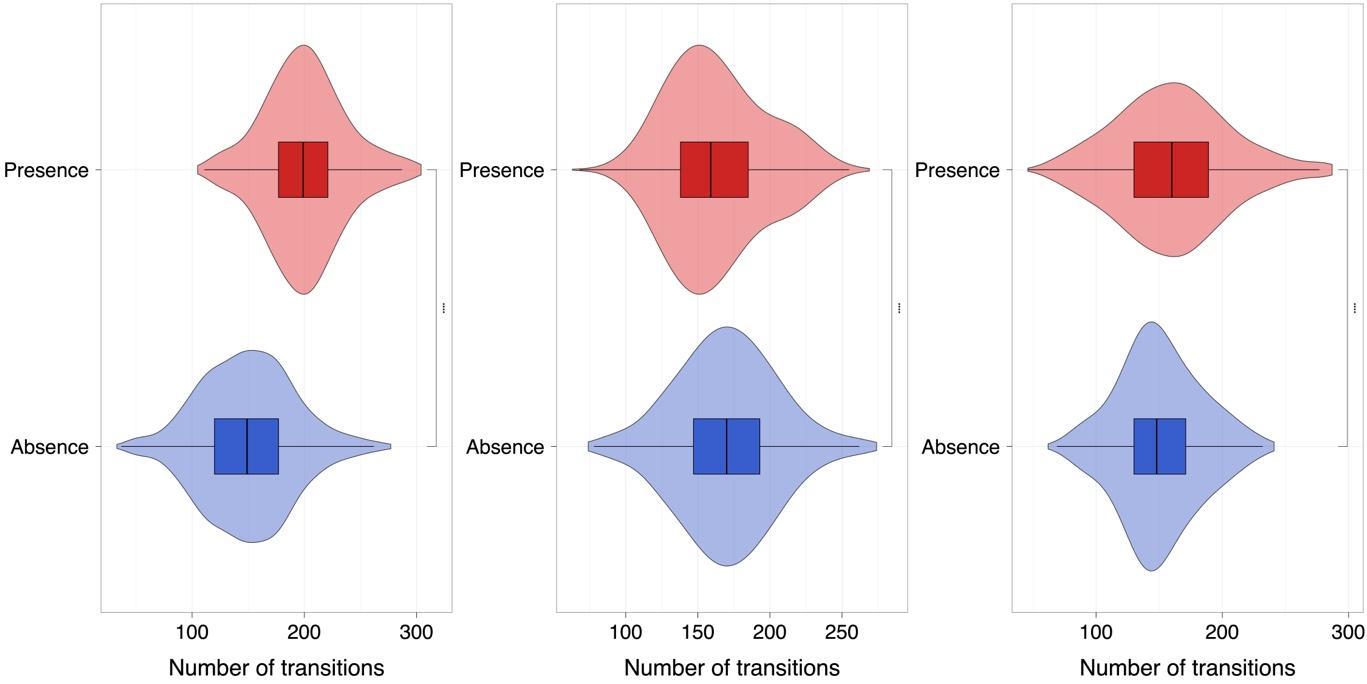

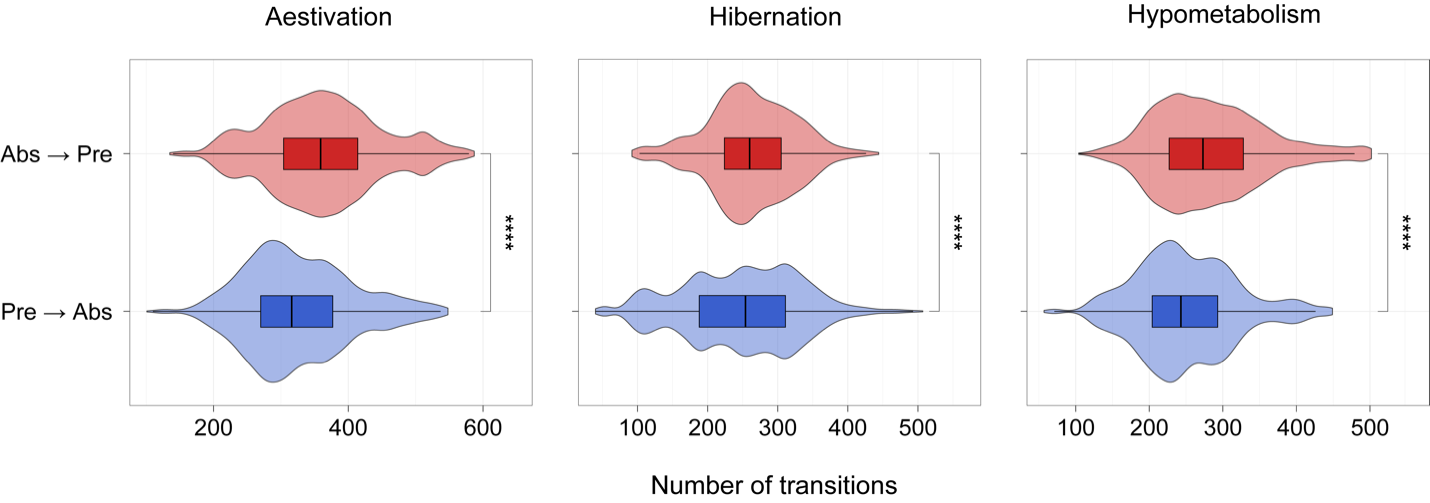

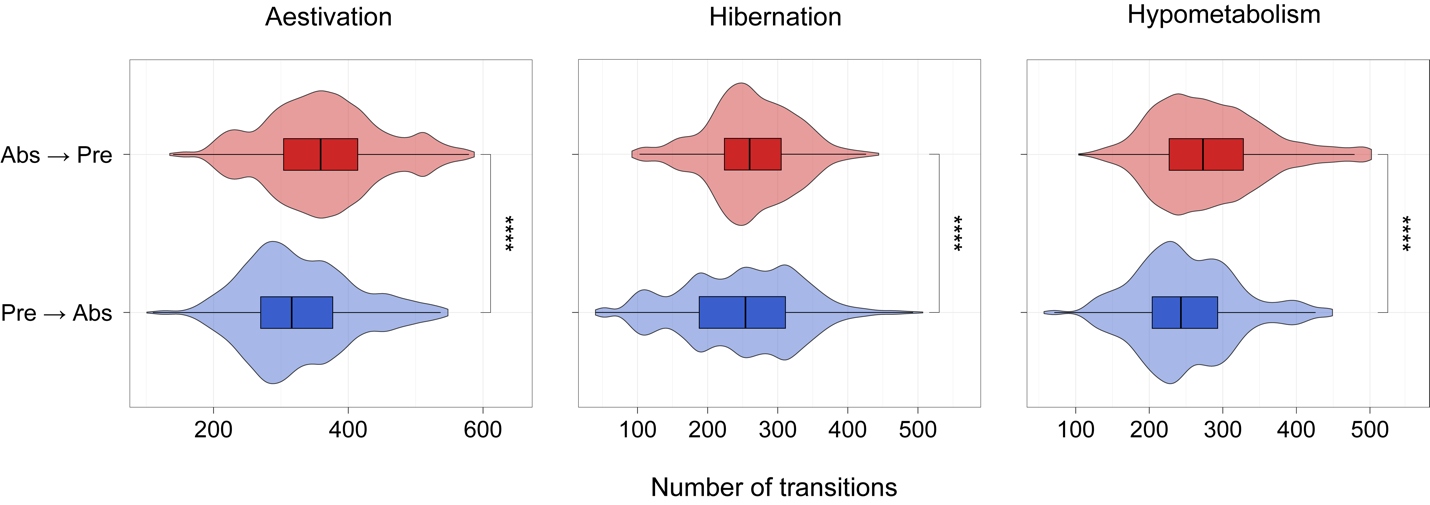

**Figure S7.** Frequency distribution for the inferred number of transitions for all the hypometabolic states for the Portik et al. (2023) phylogeny. Differences between groups were calculated using the Wilcoxon test. Significance codes for p-value are as follows (ns: > 0.05 / * ≤ 0.05 / ** ≤ 0.01 / ***: ≤ 0.001 / ****: ≤ 0.0001*).

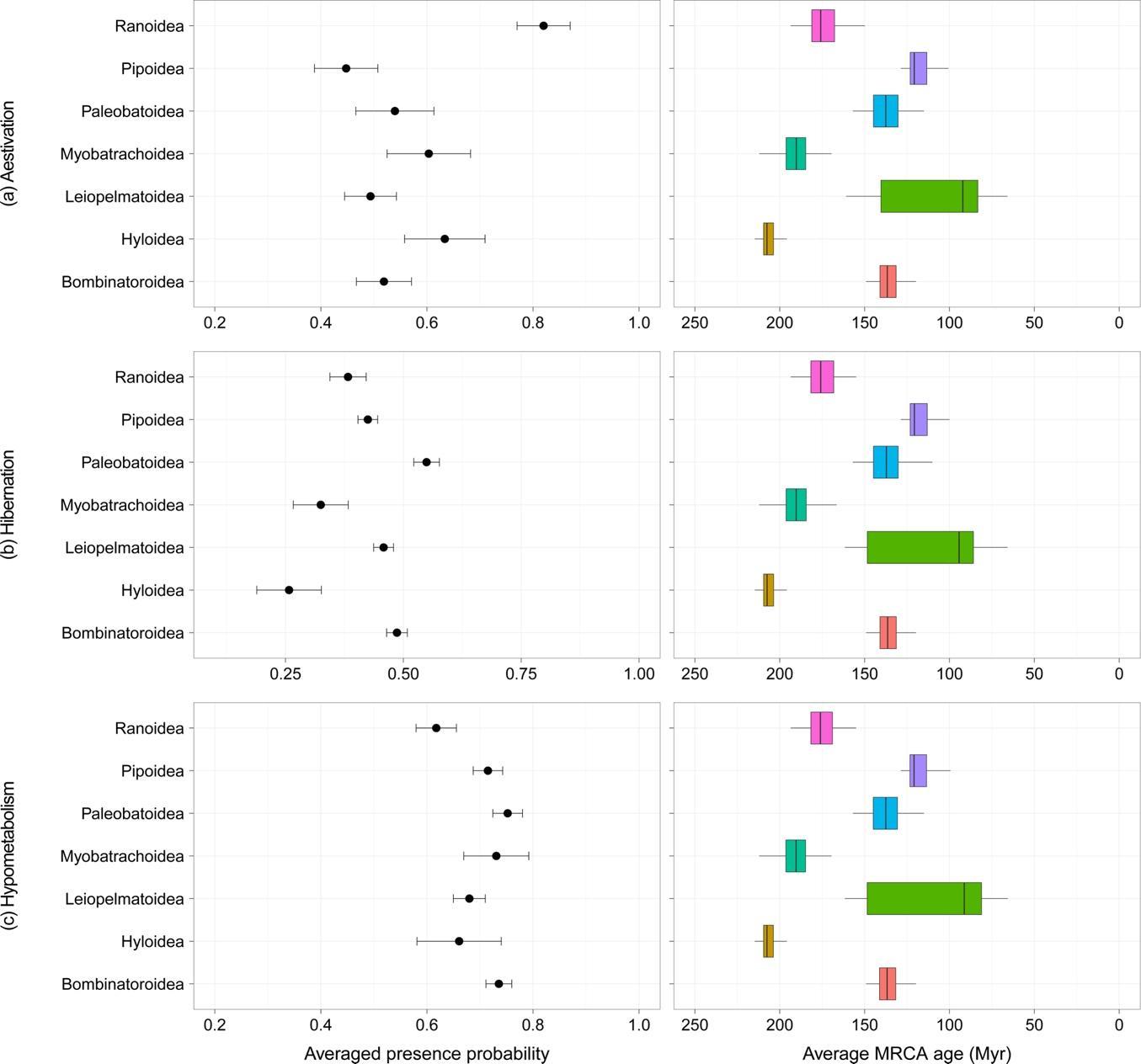

**Figure S8.** Probabilities from the ancestral state reconstruction and age node estimation with Portik et al. (2023) phylogeny for A) aestivation, B) hibernation and C) hypometabolism**.** Result from the MCMC approach to sample character histories from their posterior probability distribution of the SIMMAP analyses for 100 trees.

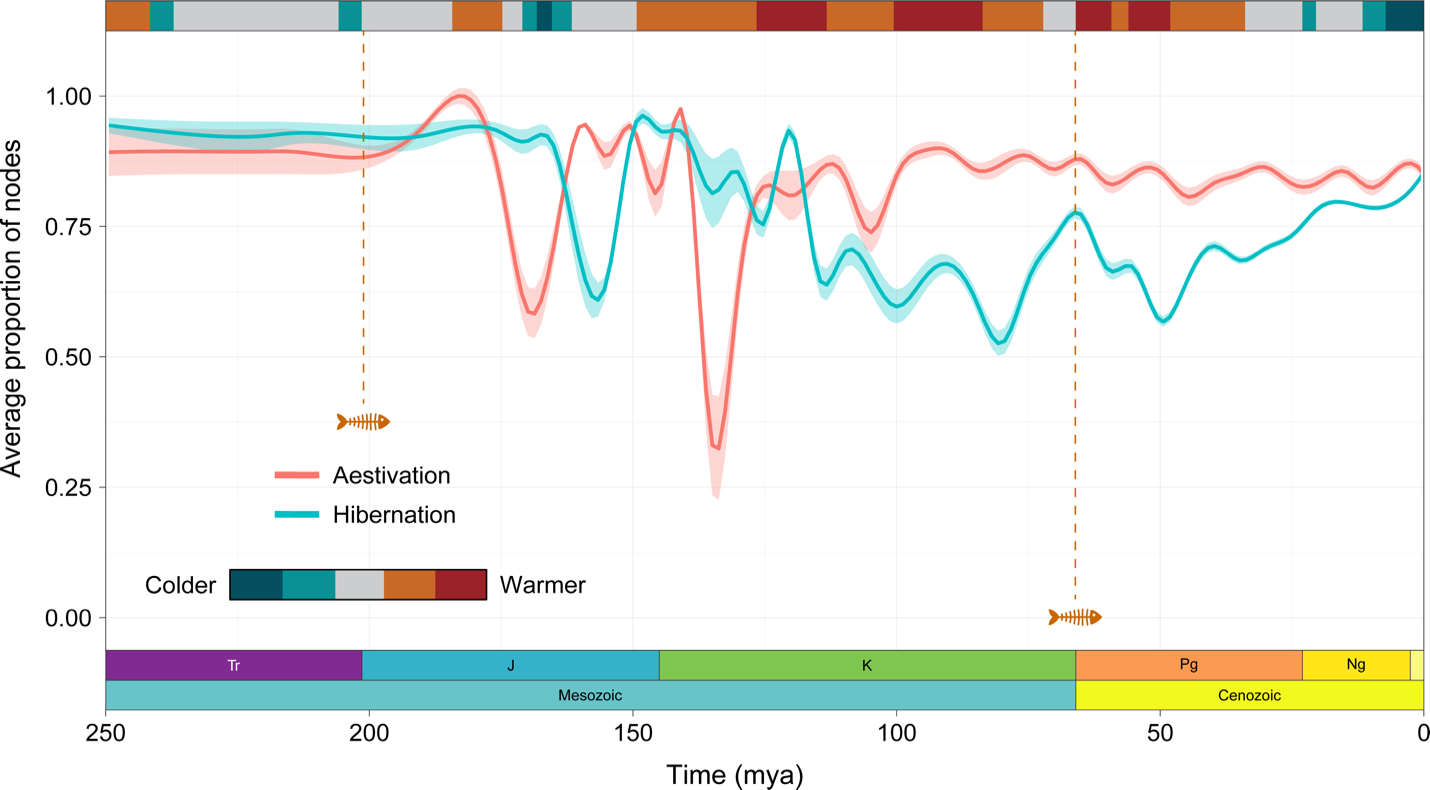

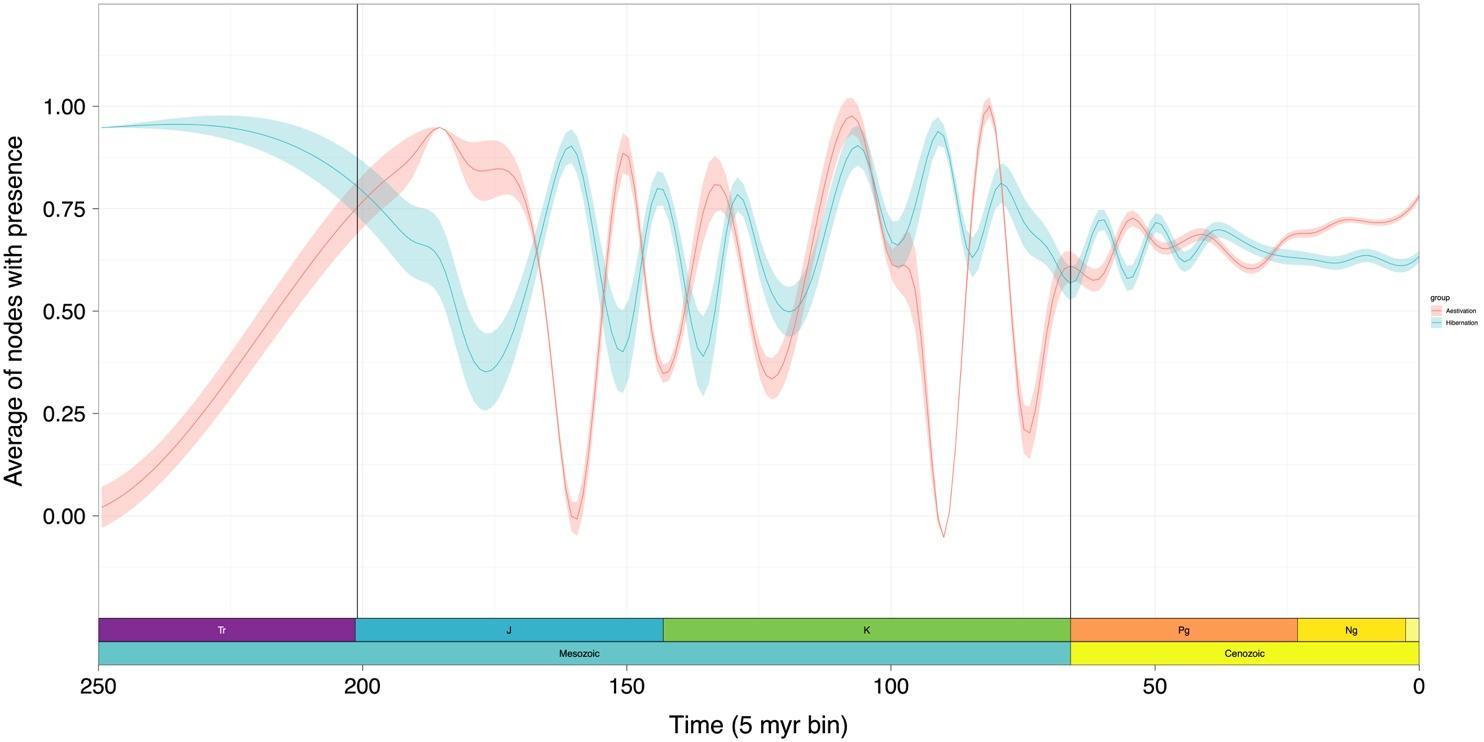

**Figure S9.** Average proportion of nodes that were inferred as presence as the most likely state for one of the two hypometabolic strategies, according to the joint ancestral state reconstruction for selected tree for Portik et al. (2023) phylogeny. The temperature bar is the same as featured in Judd et al. (2024). Curves were smoothed using the smoothing splines method and shaded areas are derived from the confidence interval (95%).

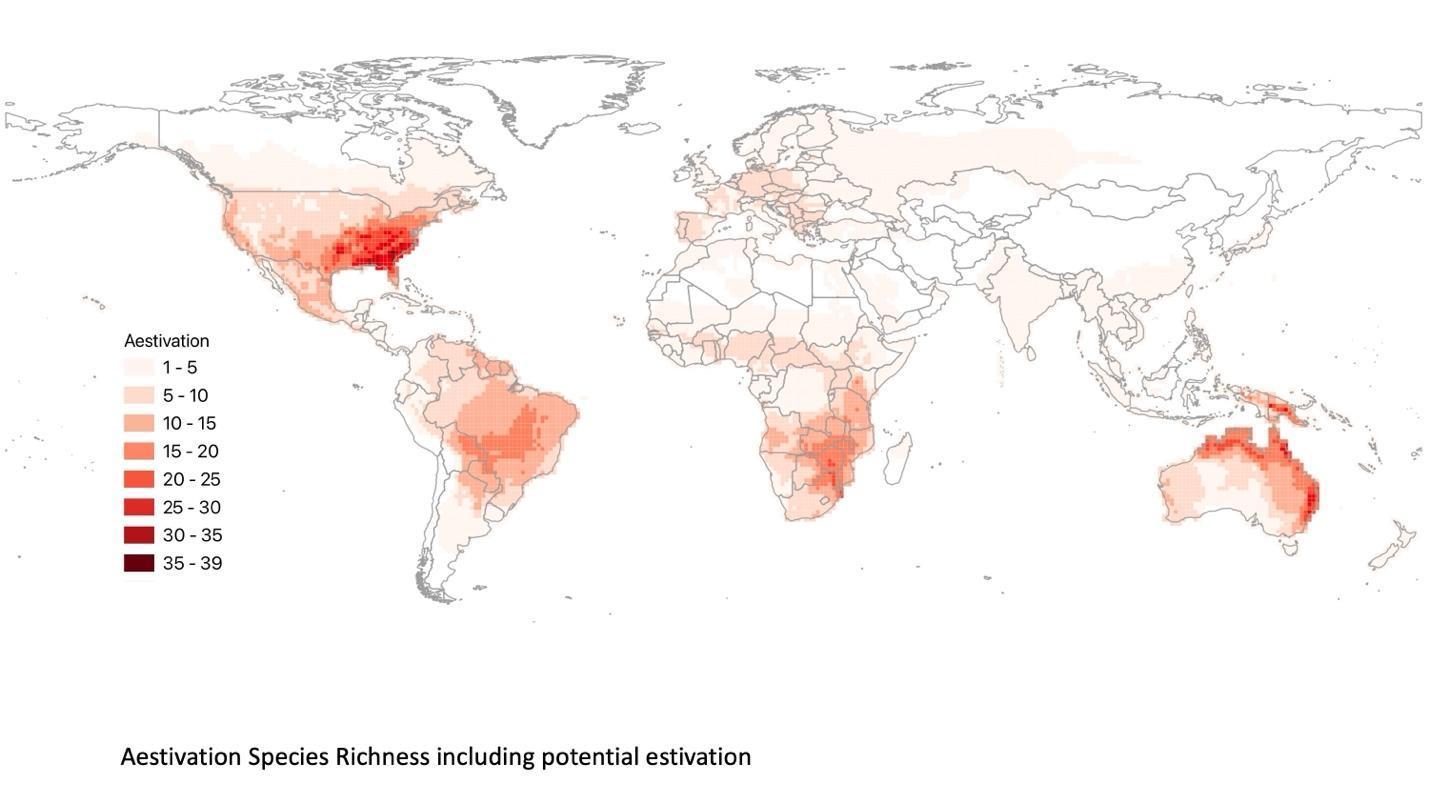

**Figure S10.** Number of species that present aestivation including potential aestivation, i.e. aestivation species richness based on the literature data.

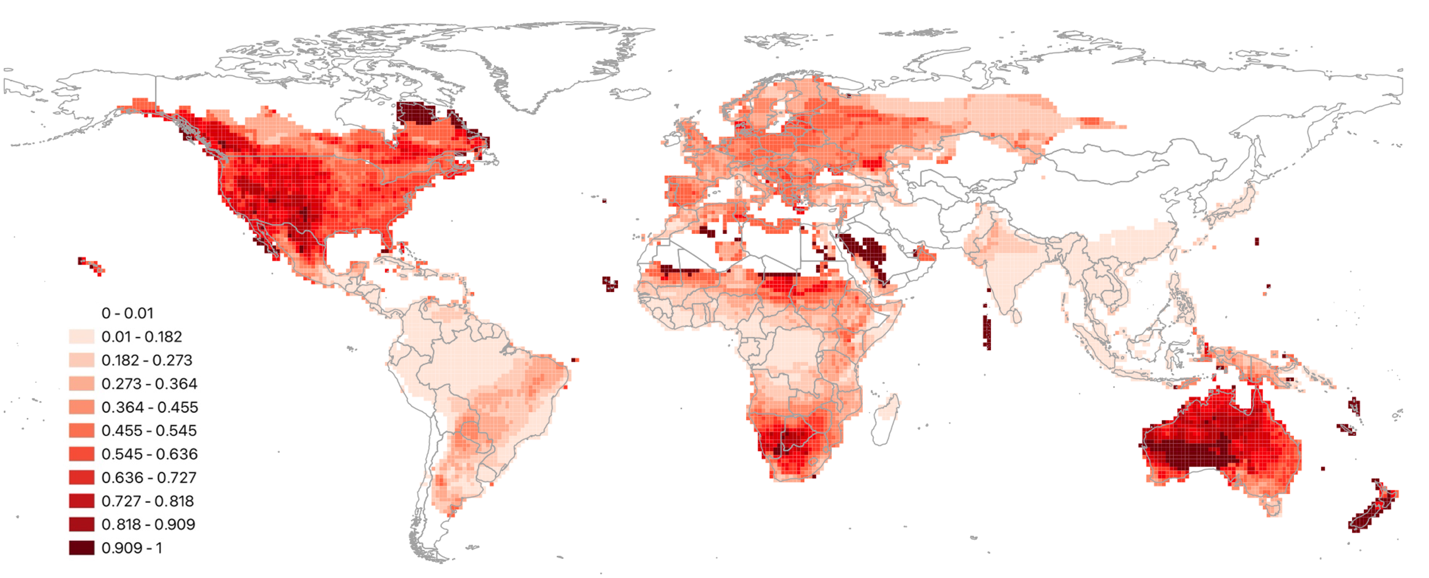

**Figure S11.** Proportion of species within the assemblage that present aestivation including potential estivation based on the literature data.

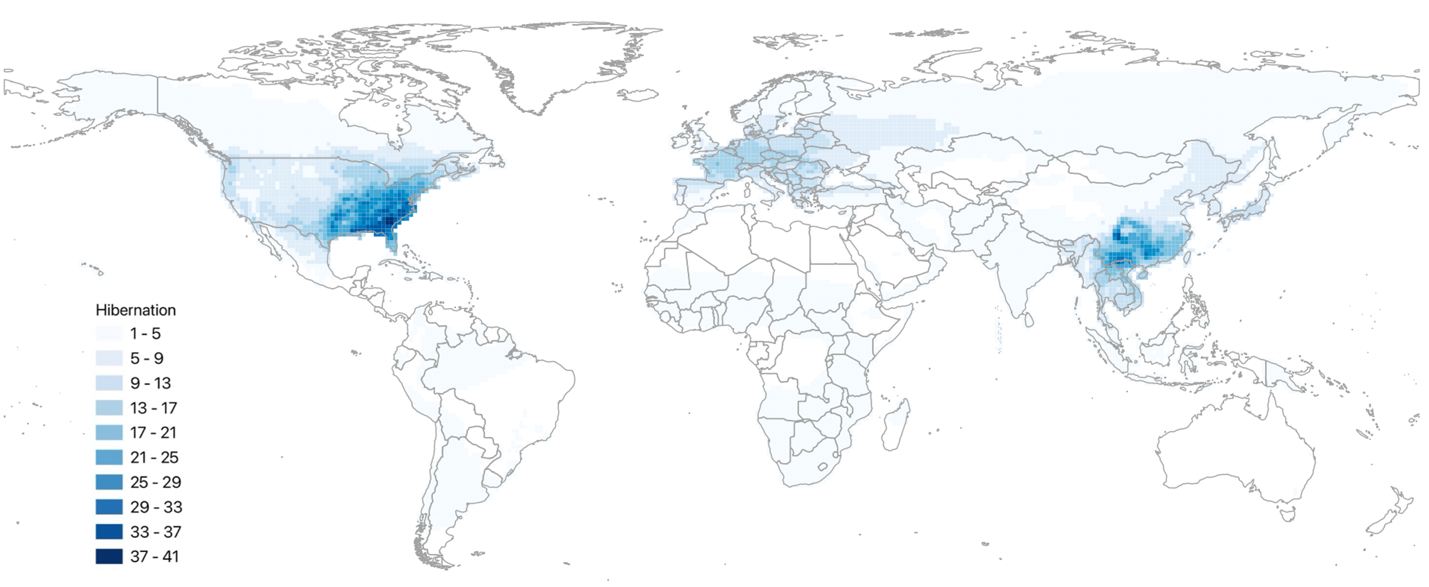

**Figure S12.** Number of species that present hibernation including potential hibernation, i.e. hibernation species richness based on the literature data. We did not include species that present cryobiosis but apparently do not present hibernation (i.e., *Litoria ewingii, Pseudacris maculata, P. streckeri,* and *Salamandrella tridactyla*).

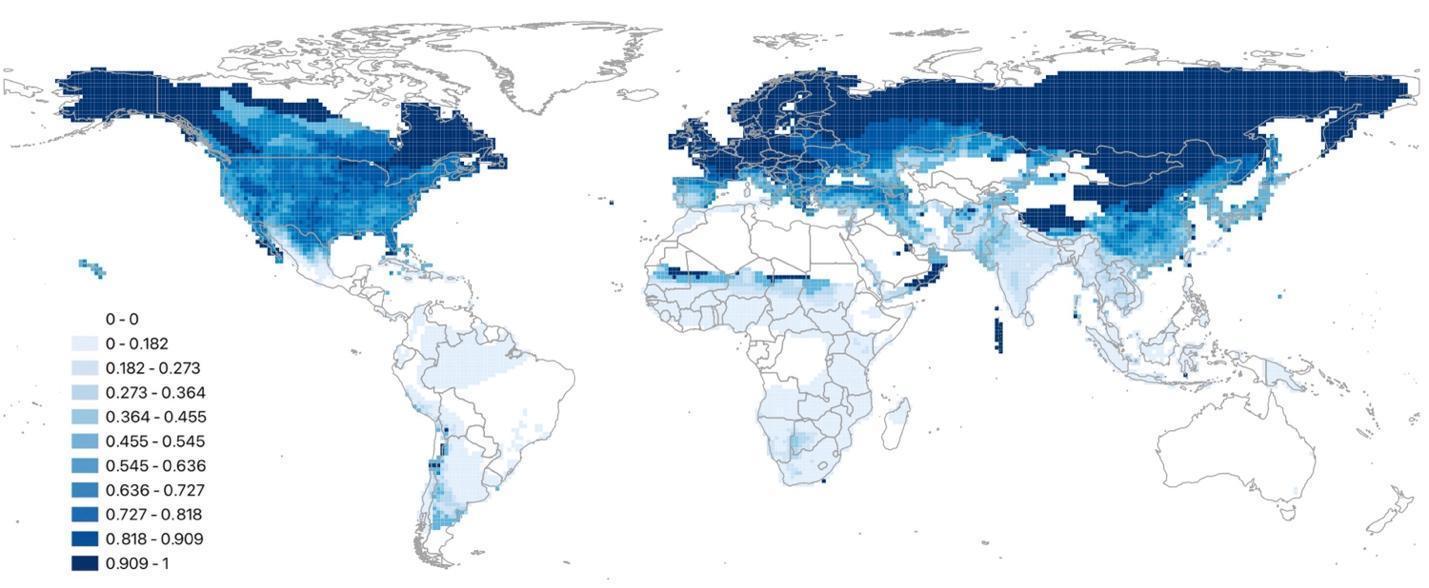

**Figure S13.** Proportion of species within the assemblage that present hibernation including potential hibernation, i.e. hibernation species richness based on the literature data. We did not include the species that present cryobiosis but apparently do not present hibernation (*Desmognathus fuscus, Litoria ewingii, Pseudacris maculata, P. streckeri,* and *Salamandrella tridactyla*).
